# Urine and Fecal Microbiota in a Canine Model of Bladder Cancer

**DOI:** 10.1101/2021.12.20.472715

**Authors:** Ryan Mrofchak, Christopher Madden, Morgan V. Evans, William C. Kisseberth, Deepika Dhawan, Deborah W. Knapp, Vanessa L. Hale

## Abstract

**Introduction:** Urothelial carcinoma (UC) is the tenth most diagnosed cancer in humans worldwide. Dogs are a robust model for invasive UC as tumor development and progression is similar in humans and dogs. Recent studies on urine microbiota in humans revealed alterations in microbial diversity and composition in individuals with UC; however, the potential role of microbiota in UC has yet to be elucidated. Dogs could be valuable models for this research, but microbial alterations in dogs with UC have not been evaluated.

**Objective:** The objective of this this pilot study was to compare the urine and fecal microbiota of dogs with UC (n = 7) and age-, sex-, and breed-matched healthy controls (n = 7).

**Methods:** DNA was extracted from mid-stream free-catch urine and fecal samples using Qiagen Bacteremia and PowerFecal kits, respectively. 16S rRNA gene sequencing was performed followed by sequence processing and analyses (QIIME 2 and R).

**Results:** Canine urine and fecal samples were dominated by taxa similar to those found in humans. Significantly decreased microbial diversity (Kruskal-Wallis: Shannon, *p* = 0.048) and altered bacterial composition were observed in the urine but not feces of dogs with UC (PERMANOVA: Unweighted UniFrac, *p* = 0.011). The relative abundances of *Fusobacterium* was also increased, although not significantly, in the urine and feces of dogs with UC.

**Conclusion:** This study characterizes urine and fecal microbiota in dogs with UC, and it provides a foundation for future work exploring host-microbe dynamics in UC carcinogenesis, prognosis, and treatment.

## 1. Introduction

Bladder cancer is the tenth most diagnosed cancer worldwide [1]. In 2020, the International Agency for Research on Cancer estimated over 573,000 new bladder cancer diagnoses would be confirmed worldwide [2]. Urothelial carcinoma (UC), also known as transitional cell carcinoma, is the most common type of bladder cancer. Age (being over age 55), race (white), sex (male), and some heritable mutations [3–10] are established risk factors for bladder cancer [11–13].

Bladder cancer is also strongly associated with environmental exposures such as smoking [14–17] or occupational exposure to chemicals like aromatic amines, pesticides, industrial dyes, or diesel fumes [18, 19]. However, not all persons exposed to these chemicals develop urothelial carcinoma indicating that there are individualized host-environment interactions that mediate UC risk.

Clear host-environment (diet) interactions mediated through the gut microbiome have emerged in colorectal carcinogenesis [20, 21] and environment-microbiome-carcinogenesis links have also begun emerging in lung cancer [22, 23]. For example, diets high in animal fat can directly or indirectly impact microbial composition by increasing liver bile acid production and excretion into the intestines. Bile tolerant microbes or microbes that can metabolize primary bile acids expand in this bile-rich environment, and some of these microbes produce pro- inflammatory, cytotoxic, or genotoxic secondary metabolites that can contribute to colorectal carcinogenesis. Work on the gut microbiome has far outpaced and outnumbered studies on the urine / bladder microbiome; however, it has now become apparent that the urine microbiota play a key role in host health and may also be influencing bladder cancer development and progression [24]. Alterations in urine microbiota have been reported in association with multiple genitourinary diseases including chronic kidney disease [25], chronic prostatitis, chronic pelvic pain syndrome [26], interstitial cystitis [27], sexually transmitted infections [28], urgency urinary incontinence [29], urinary tract infections [30], urinary stone disease [31], urogenital schistosomiasis [32], urogynecologic surgery [33], and vaginosis [34]. A few recent studies on the urine / bladder microbiome have also revealed subtle but intriguing differences in urine or bladder tissue microbial diversity and composition of individuals with and without UC (**Table 1**) [17,35–45], but approaches and results in these studies vary widely. Studies in relevant animal models could advance this research by offering a more controlled environment. Multiple animal models of UC have been described, with most being rodent models that have many limitations [46].

**Table 1:**
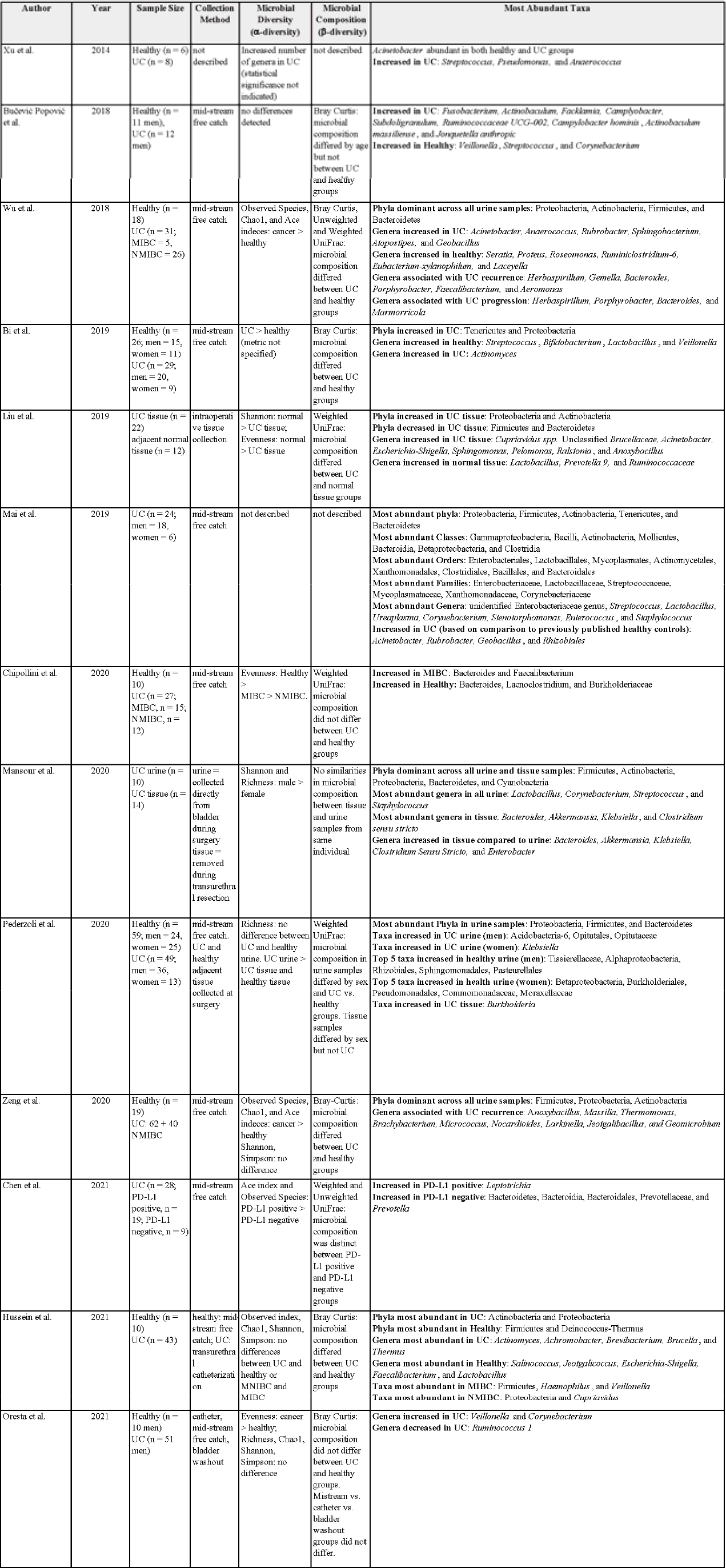
Key findings in 13 publications about the urine / tissue microbiota and urothelial carcinoma. . MIBC = Muscle Invasive Bladder Cancer; NMIBC = Non-Muscle Invasive Bladder Cancer; PD-L1 = Programmed Cell Death 1 Ligand 1; UC = Urothelial Carcinoma.

The focus of this study was on invasive UC utilizing a naturally-occurring canine model and comparing the urine and fecal microbiota of dogs with and without UC. While it can be difficult to produce the collective features of cancer heterogeneity, molecular features, aggressive cancer behavior, and host immunocompetence in experimental models, these features are present in the canine model [57–59]. In humans, approximately 25 % of all UC cases are muscle invasive [44] while in dogs with UC, over 90 % present with intermediate- to high-grade muscle invasive bladder cancer [47, 48]. Moreover, humans and dogs share many of the same environmental exposures, and canine UC, like human UC, has been epidemiologically linked to chemical exposures including herbicides and pesticides [49, 50]. Dogs also exhibit strong heritable (breed-specific) associations with UC offering unique opportunities for gene-environment studies [49–51]. Notably, the human microbiome is more similar to the dog microbiome compared to other animal models, such as the rodent microbiome [52], making dogs a more suitable model for studying microbiota in relation to UC.

## 2. Materials and Methods

### 2.1 Sample Collection

All dogs were recruited through Purdue University College of Veterinary Medicine between September 2016 and October 2019 (Purdue IACUC: 1111000169; Ohio State University IACUC: 2019A00000005). Urine and fecal samples were initially collected from 57 dogs with biopsy-confirmed urothelial carcinoma (UC) and 56 age, sex, and breed-matched healthy controls (**Figure 1**). Dogs with active urinary tract infections were excluded. We additionally excluded any dog with a history of chemotherapy (vinblastine, zebularine, vemurafenib, chlorambucil, mitoxantrone, and cyclophosphamide) or a history of antibiotics within the previous 3 weeks due to the potential effects of these medications on the microbiome [53–60]. We did not exclude dogs on non-steroidal anti-inflammatory drugs (NSAIDs), including piroxicam and deracoxib, which are commonly used in dogs with UC. Healthy dogs underwent physical exams and had no history of antibiotics (within the previous 3 weeks) or indications of gastrointestinal or urogenital disease.

**Figure 1:**
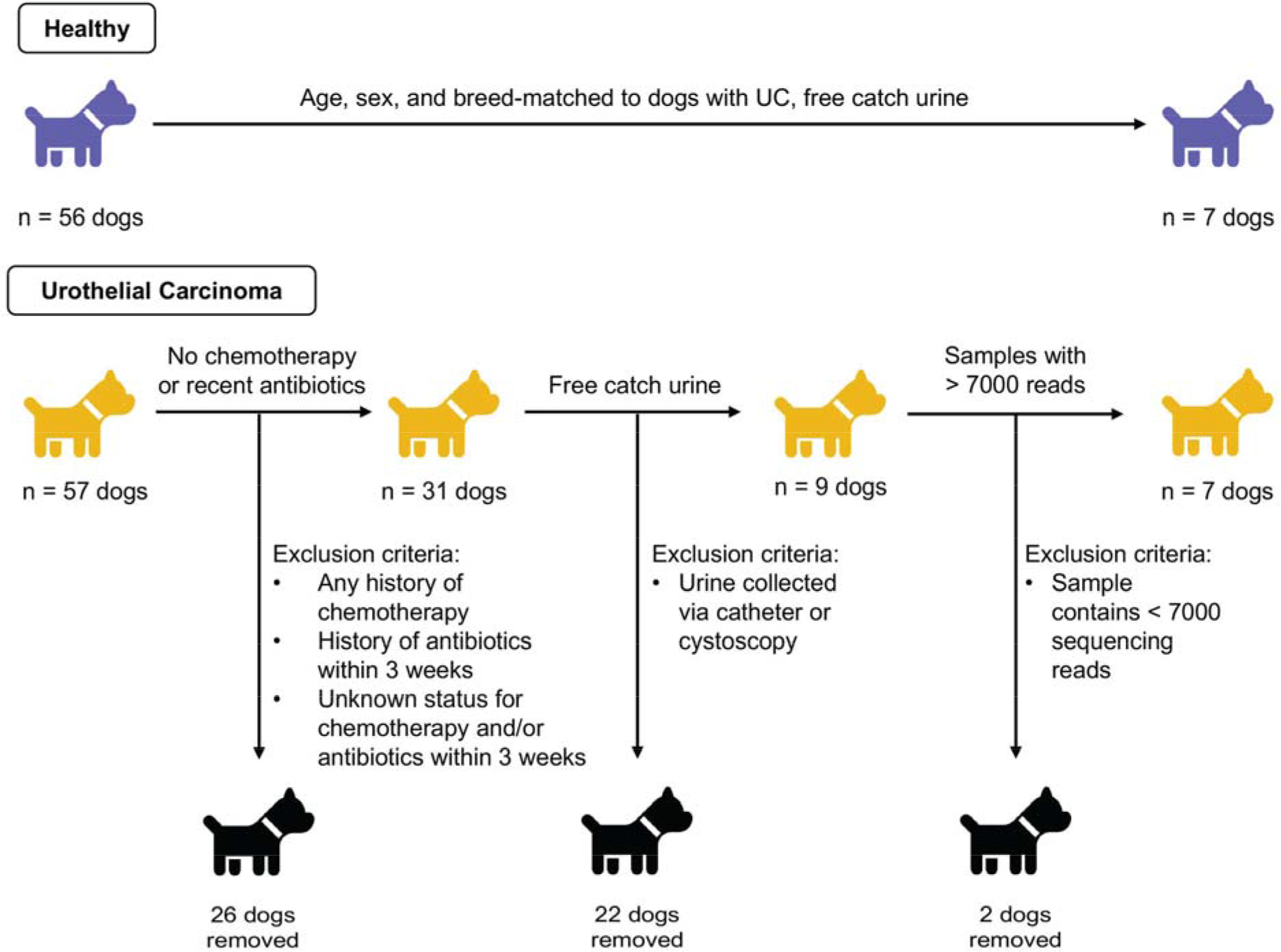
Experimental design.

In healthy dogs, urine was collected via mid-stream free catch. In dogs with UC, a variety of urine collection methods were employed as deemed clinically appropriate including: mid- stream free catch, catheter, or cystoscopy. Free catch urine can include bacteria from the bladder, urethra, periurethral skin, prepuce, or vagina, while urine collected via catheterization or cystoscopy primarily includes microbes from the bladder and limits the presence of genital and skin microbes [41,61–63]. To determine if collection method could potentially influence our results, we compared samples from dogs with UC collected via free catch (n = 8) to samples collected via non-free catch methods (catheterization, cystoscopy) (n = 11) (**Supp. Table 1; Supp. Figures 1,2,3**). We observed significant differences in microbial composition but not diversity by collection method (Bray-Curtis PERMANOVA rarefied: *p* = 0.008; non-rarefied: *p* = 0.005; **Supp. Figures, 1f,2f**). Moreover, *Staphylococcus* and *Streptococcus* – common skin colonizers - were amongst the top genera in free catch urine but not amongst the top genera in non-free catch urine (**Supp. Table 2**). Based on the compositional differences we observed by collection method and on other studies that have reported differences in urine microbiota due to collection method [41,61–65], we opted to limit the remainder of our analyses to samples collected via free catch only. This allowed us to compare microbiota in urine from healthy dogs and dogs with UC without introducing collection method as a potential confounder.

**Figure 2:**
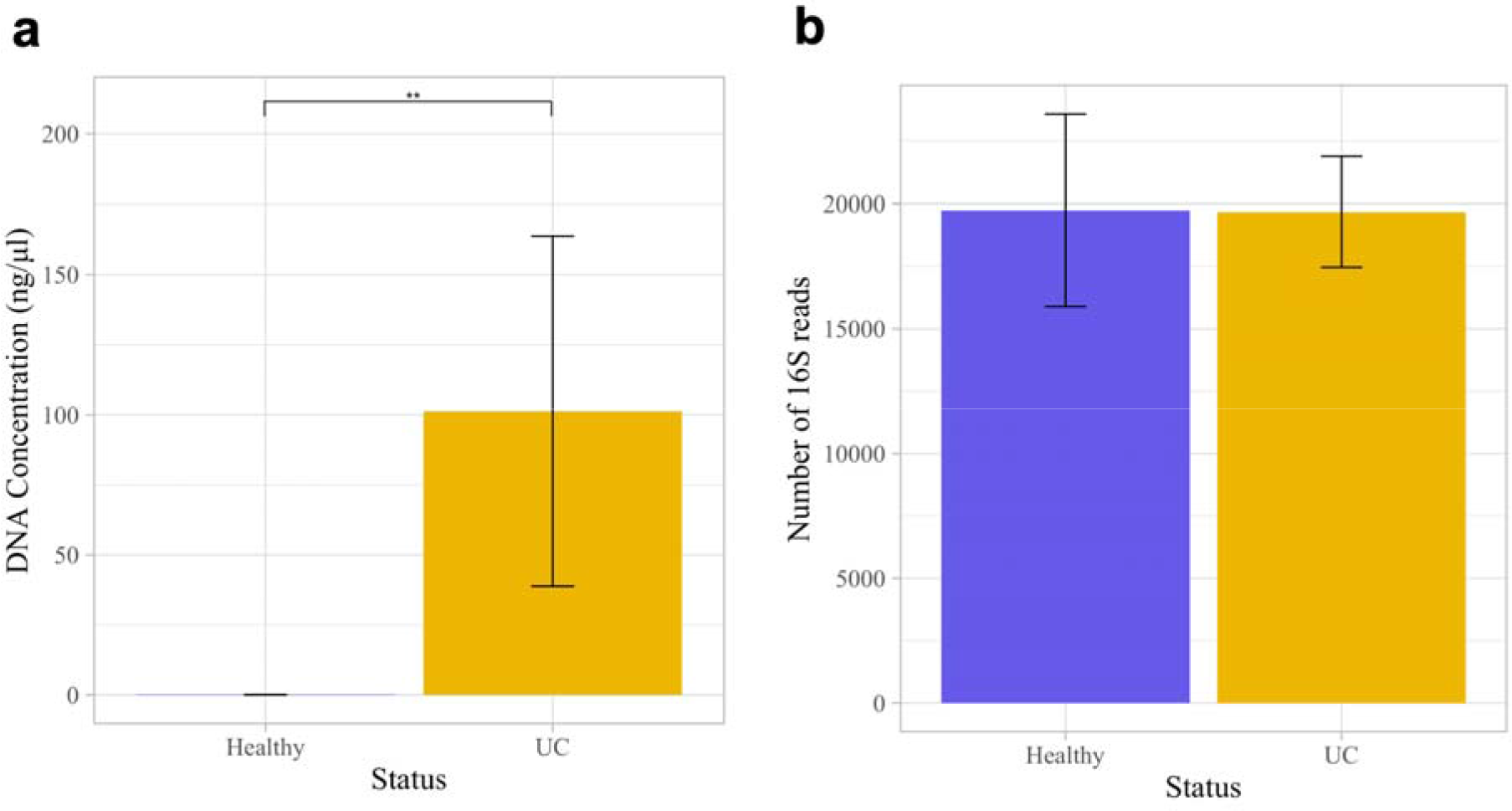
DNA concentrations and number of 16S reads in the urine samples of dogs with and without urothelial carcinoma (UC). (**a**) DNA concentrations were significantly greater in dogs with UC than in healthy dogs (Wilcoxon Rank Sum test, *p* = 0.002). (**b**) The number of 16S reads did not differ significantly between groups (two-sample t-test, *p* = 0.99). Error bars denote standard error. Statistical significance is represented by stars: * < 0.05, ** < 0.001, *** < 0.0001

**Table 2:** Demographics of dogs with and without urothelial carcinoma (UC). Urine samples were collected and analyzed from all dogs. Stool samples were collected and analyzed from a subset of these dogs including 6 healthy (4 females, 2 males), and 4 with UC (3 females, 1 male).

| Category | Healthy | UC |
| --- | --- | --- |
| Sex, n (%) |  |  |
| Females | 5 (71.4 %) | 5 (71.4 %) |
| spayed | 4 | 4 |
| non-spayed | 1 | 1 |
| Males | 2 (28.6 %) | 2 (28.6 %) |
| neutered | 2 | 2 |
| non-neutered | 0 | 0 |
| Age (mean ± SD) | 10.1 ± 1 | 10.1 ± 0.7 |

As such, after exclusions, urine samples from a total 7 dogs with UC and 7 age, sex, and breed-matched healthy controls were compared in this study (**Table 2)**. Fecal microbiota from a subset of these 14 dogs for which we had fecal samples (4 dogs with UC and 6 healthy controls) were also compared [30,66,67]. All urine and stool samples were placed on ice immediately after collection and then transferred into a -80°C freezer. Samples were transported on dry ice from Purdue (West Lafayette, IN, USA) to the Ohio State University (Columbus, OH, USA), where they were stored in at -80°C until extraction.

### 2.2 DNA extraction and quantification

Urine samples were extracted using QIAamp^®^ BiOstic^®^ Bacteremia DNA Isolation Kit (Qiagen, Hilden, Germany) as described previously [68]. Fecal samples were extracted using the QIAamp^®^ PowerFecal^®^ DNA Kit (Qiagen, Hilden, Germany) following the manufacturer’s instructions. Negative (no sample) controls were run with each kit used for extraction. DNA concentrations were measured using a Qubit^®^ 4.0 Fluorometer (Invitrogen, Thermo Fisher Scientific^TM^, Carlsbad, CA, USA) and purity was assessed using Nanodrop One (Thermo Fisher Scientific^TM^, Carlsbad, CA, USA).

### 2.3 16S rRNA sequencing and sequence processing

Library preparation, PCR amplification, and amplicon sequencing was performed at Argonne National Laboratory (DuPage County, Illinois). Likewise, negative controls underwent the full extraction, library preparation, and sequencing process. We amplified the V4 region of the 16S rRNA gene using primers 515F and 806R, and PCR and sequencing were performed as described previously (2 x 250bp paired-end reads, on an Illumina Miseq (Lemont, IL, USA)) [68–70]. Raw, paired-end sequence reads were processed using QIIME2 v. 2020.11 and DADA2 [71, 72]. Taxonomy was assigned in QIIME2 using the Silva 132 99% database and the 515F / 806R classifier [73, 74]. In the analysis comparing urine collection method in dogs with UC, we excluded samples with fewer than 1,000 reads and analyzed the data with rarefaction (at 1,000 reads) and without rarefaction. We included both analyses because rarefaction, especially at low read counts, can increase type 1 errors and mask potential differentially abundant taxa between samples [75]. In the analyses comparing urine and fecal microbiota from dogs with and without UC, samples with fewer than 7,000 reads were excluded; this cutoff allowed us to retain all but two urine samples while excluding all negative controls (**Figure 1**). Urine samples from dogs with and without UC were rarefied at 7,000 reads; fecal samples were rarefied at 9,233 reads, which included all fecal samples. Sequencing data for this project is available in SRA BioProject PRJNA76392.

### 2.4 Urine and fecal sequence data processing

Prior to analyses, we first removed singletons (Amplicon Sequence Variants (ASVs) with only one read in the dataset). ASVs are roughly equivalent to a microbial species or strain. We then applied the R package decontam to identify and filter out putative contaminant ASVs based on their frequency and prevalence (0.5 threshold) as compared to negative controls (R package, v.1.10.0) [76]. In total, we identified and removed 13 putative contaminant ASVs from the urine samples and 8 from the fecal samples (**Supp. Table 3**). We also removed sequences aligned to chloroplasts, eukaryotes, mammalia, and mitochondria. In addition, in the urine samples, we removed taxa within the phylum Cyanobacteria and the class Chloroflexia. All six negative controls, which contained fewer than 7000 reads, were then removed from subsequent analyses.

### 2.5. Statistical analyses

Data were tested for normality using the Shapiro Wilk Normality Test in R version 3.5.2 [77]. We then compared DNA concentrations and read numbers between groups using Wilcoxon Rank Sum tests and two-sample t-tests, respectively. All alpha and beta diversity metrics were assessed using the R package phyloseq with a p-value cutoff of 0.05 adjusted using the Benjamini & Hochberg False Discovery Rates [78]. Alpha-diversity metrics included Shannon, Simpson, and Observed Features followed by Kruskal-Wallis Rank Sum Tests to compare metrics by group. Beta-diversity metrics included Bray-Curtis, Unweighted UniFrac, and Weighted UniFrac. Permutational Multivariate Analysis of Variance (PERMANOVA) were implemented in QIIME2 v. 2020.11 to compare bacterial community composition by group. An Analysis of Composition of Microbiome (ANCOM) was used to identify differentially abundant taxa by group.

## 3. Results

### 3.1 Urine microbiota in dogs with UC

We compared the urine microbiota of 7 dogs with UC to 7 age, sex, and breed-matched healthy controls. The total number of reads across all samples ranged from 7,232 – 36,692 with a mean of 20,010 ± 7,329 reads. Urine samples contained a total of 21 bacterial phyla, 308 genera, and 187 species. Urine DNA concentrations were significantly higher in dogs with UC as compared to healthy dogs (**Figure 2a**: Wilcoxon Rank Sum test, *p* = 0.002), but there was no significant difference in the number of 16S reads between dogs with and without UC (**Figure 2b**: two-sample t-test, *p* = 0.99).

Dogs with UC had significantly lower urine microbial diversity compared to healthy dogs as measured by the Shannon diversity index and Observed Features but not by the Simpson diversity index (Kruskal-Wallis: Shannon, *p* = 0.048; Observed Features, *p* = 0.025; Simpson, *p* = 0.133; **Figure 3a, Supp. Figure 4a,b**). Dogs with UC also had significantly different urine microbial composition than healthy dogs based on an Unweighted UniFrac distance matrix (**Figure 3b**; PERMANOVA, *p* = 0.011); although, no significant differences were observed by Bray Curtis (*p* = 0.888) or Weighted UniFrac (*p* = 0.168) distance matrices (**Supp. Figure 4c,d**). At the phylum level, Firmicutes (healthy: 61.1 %; UC: 79.5 %) Proteobacteria (healthy: 18.0 %; UC: 15.6 %), and Actinobacteria (healthy: 12.5 %; UC: 4.26 %) were the three most abundant phyla in the urine of healthy dogs and dogs with UC (**Figure 4a**). At the family level, Staphylococcaceae (healthy 42.6%; UC 48.6%) and Streptococcaceae (healthy 5.99 %; UC 14.8%) were amongst the most abundant taxa (**Figure 4b;** For genus and order level taxa see **Supp. Figure 5)**. Interestingly, *Fusobacterium* was present in the urine of dogs with UC but not in the urine of healthy dogs (relative abundance of *Fusobacterium* in healthy dogs: 0 %; in dogs with UC: 0.167 %). There were no differentially abundant taxa between healthy dogs and dogs with UC at the phylum, genus, or ASV levels.

**Figure 3:**
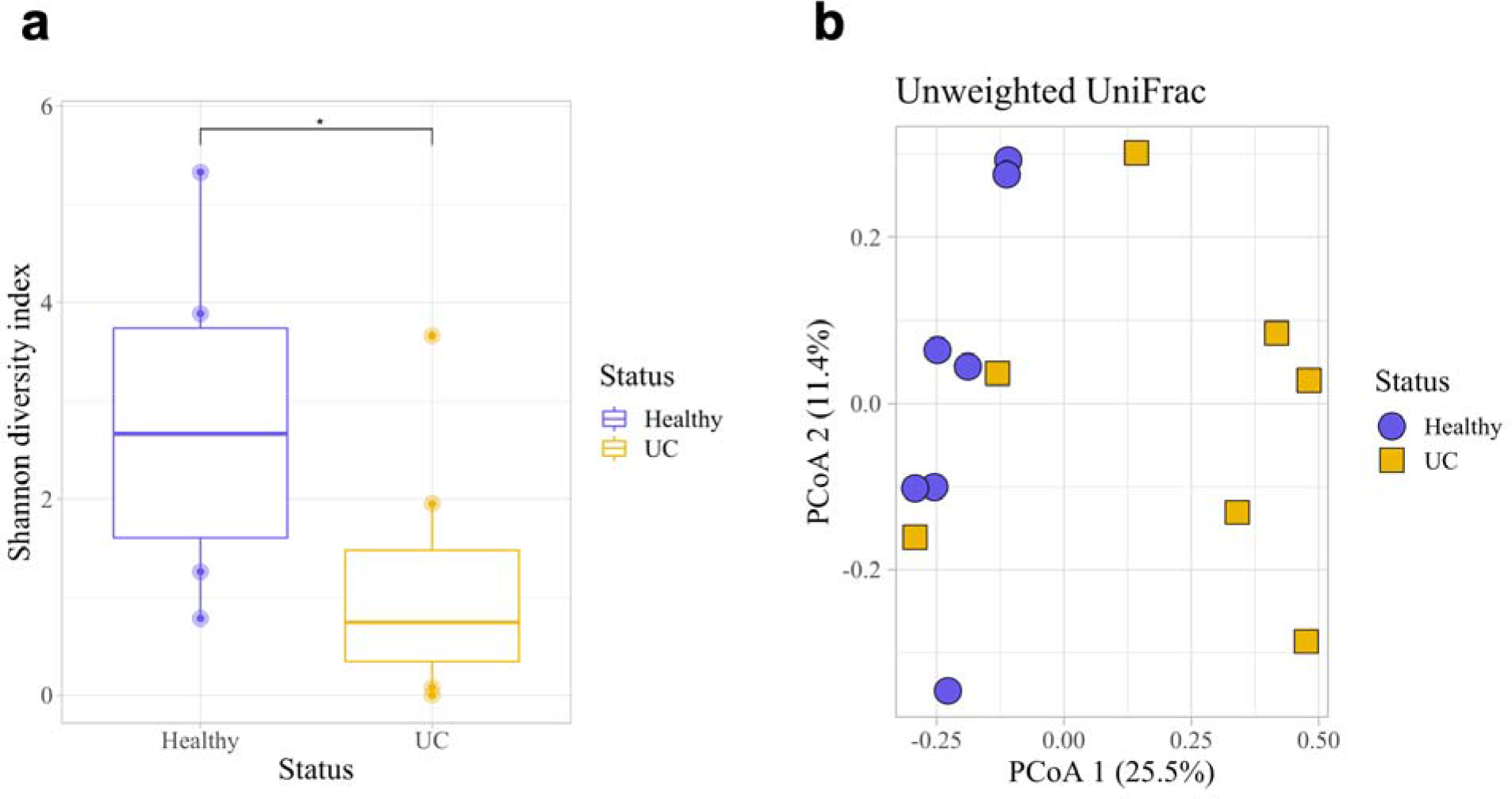
Microbial diversity and composition in the urine of dogs with and without UC. (**a**) Healthy dogs had a significantly higher microbial diversity compared to dogs with UC as measured by the Shannon diversity index (Kruskal-Wallis, *p* = 0.048). (**b**) Microbial composition between healthy dogs and dogs with UC also differed significantly (Unweighted UniFrac, PERMANOVA, *p* = 0.011). Error bars denote standard error. Statistical significance is represented by stars: * < 0.05, ** < 0.001, *** < 0.0001

**Figure 4:**
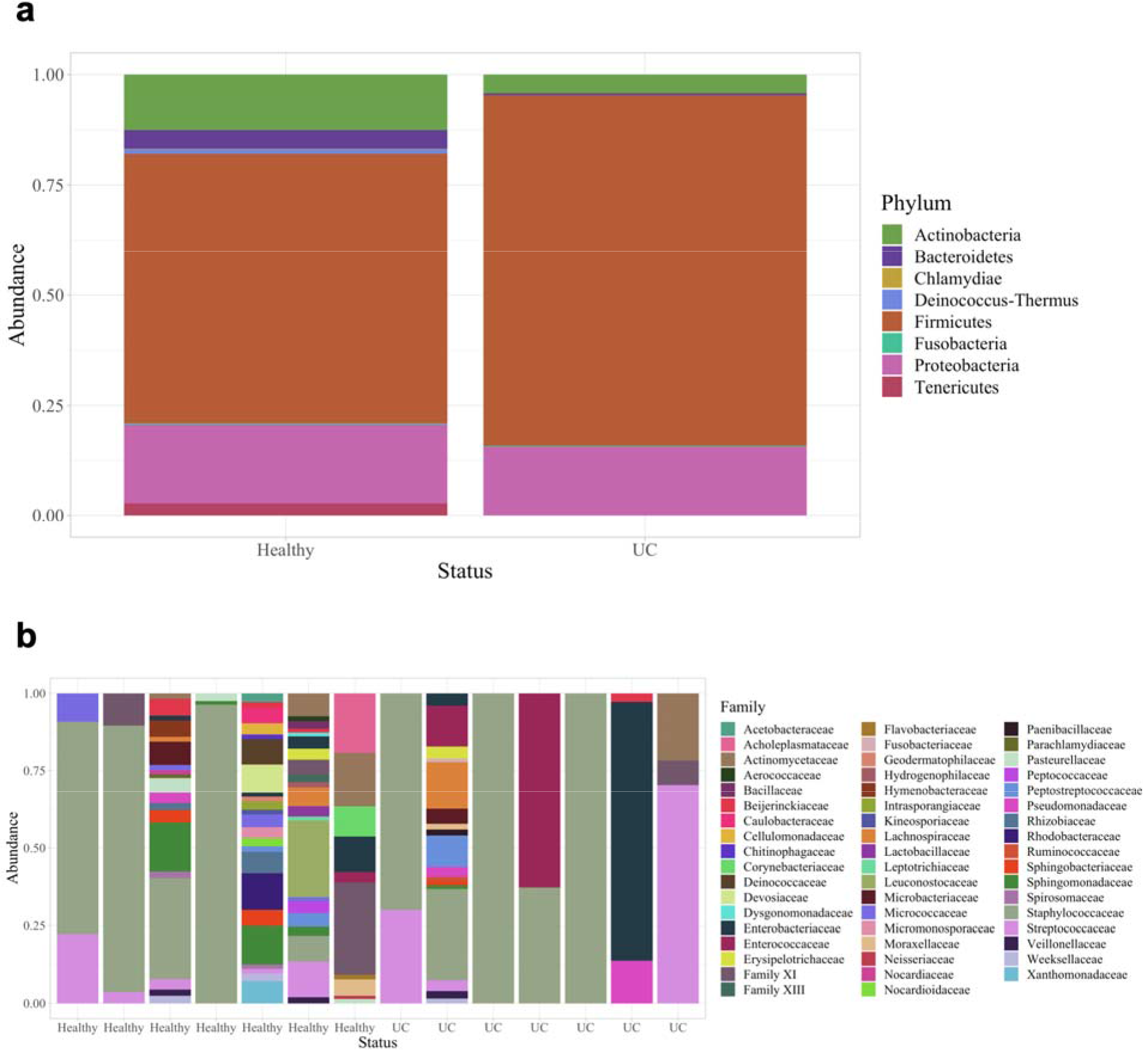
Phyla and family taxa bar plots of urine samples in dogs with and without UC. (**a**) Phyla and (**b**) family relative abundances. At the family level, the taxonomic composition of each sample is shown individually to demonstrate the variability across urine samples.

**Figure 5:**
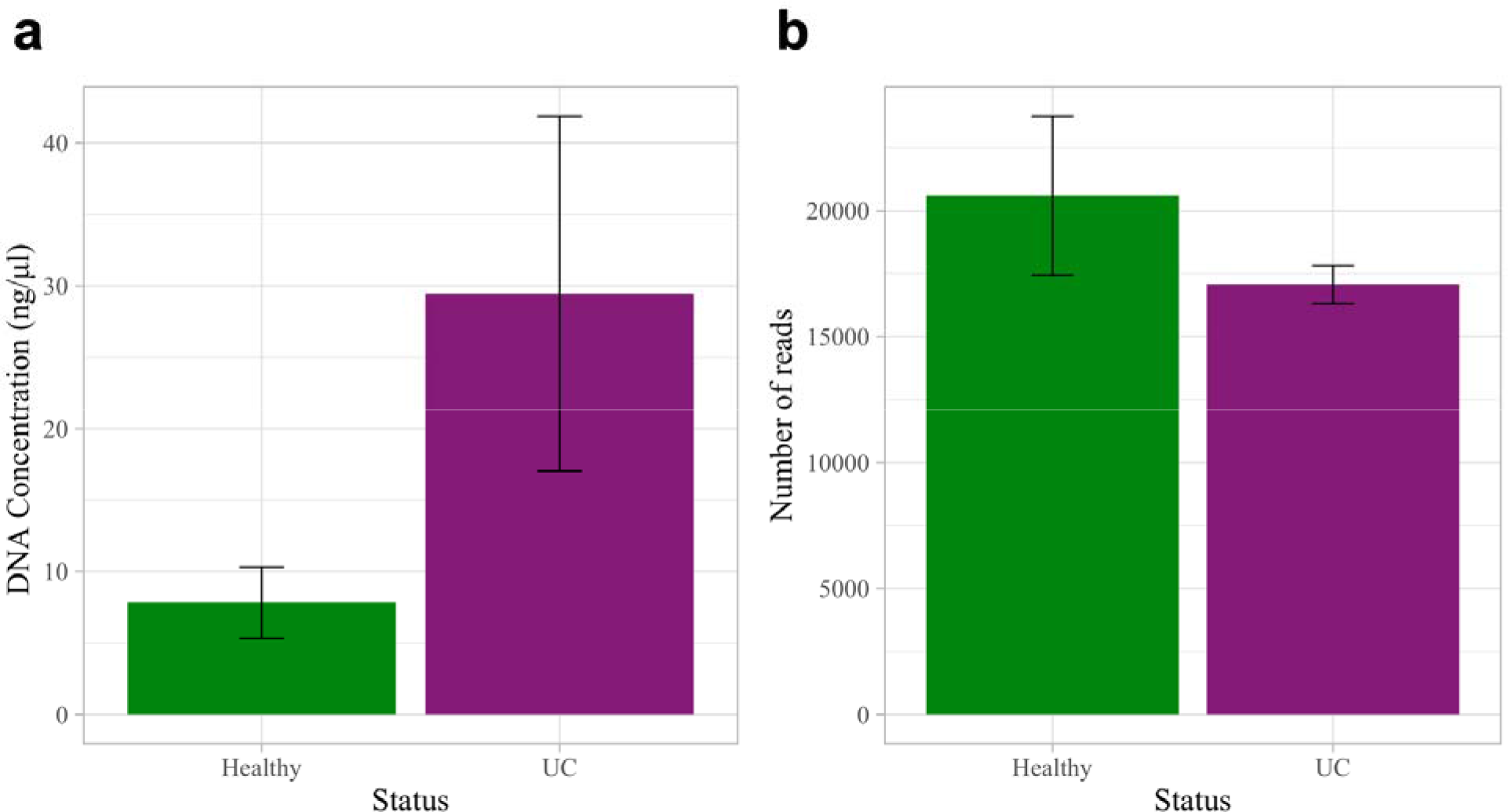
DNA concentrations and number of 16S reads in the fecal samples of dogs with and without UC. (**a**) DNA concentrations were greater (but not significantly) in dogs with UC as compared to healthy dogs (Wilcoxon Rank Sum Test, *p* = 0.136). (**b**) The number of 16S reads did not differ significantly between groups (two-sample t-test, *p* = 0.322). Error bars denote standard error.

### 3.2 Fecal microbiota in dogs with UC

We compared the fecal microbiota of a subset of dogs from the urine analyses for which we also had fecal samples: four dogs with and six dogs without UC. The total number of reads across all fecal samples ranged from 9,233 – 28,345 with a mean of 19,196 ± 6,100 reads. Fecal samples contained a total of 8 bacterial phyla, 92 genera, and 45 species. There was no significant difference in fecal DNA concentrations or number of 16S reads in dogs with UC as compared to healthy dogs; although, DNA concentrations were greater in dogs with UC (DNA concentration: Wilcoxon Rank Sum Test, *p* = 0.136; 16S reads: Two-sample t-test, *p* = 0.322; **Figure 5**).

Fecal microbial diversity and composition did not differ significantly in dogs with and without UC (Kruskal-Wallis: Shannon, *p* = 0.67; Unweighted UniFrac PERMANOVA, *p* = 0.252; **Figure 6**, **Supp. Figure 6**). The top three most abundant phyla across all fecal samples were Firmicutes (healthy: 72.6 %; UC: 32.9 %), Bacteroidetes (healthy: 10.6 %, UC 31.9 %) and Fusobacteria (healthy: 11.3 %, UC: 31.1 %) (**Figure 7; Supp. Figure 7**). At the family and genera levels, Fusobacterieacea (healthy: 11.4 %, UC: 31.7 %) and *Fusobacterium* (healthy: 12.0 %, UC: 33.1 %) were the most abundant taxa in UC but not healthy samples, respectively; although, these differences were not statistically significant. Only one *Bacteroides spp*. was significantly increased in relative abundance in dogs with UC compared to healthy dogs (ANCOM, W = 25).

**Figure 6:**
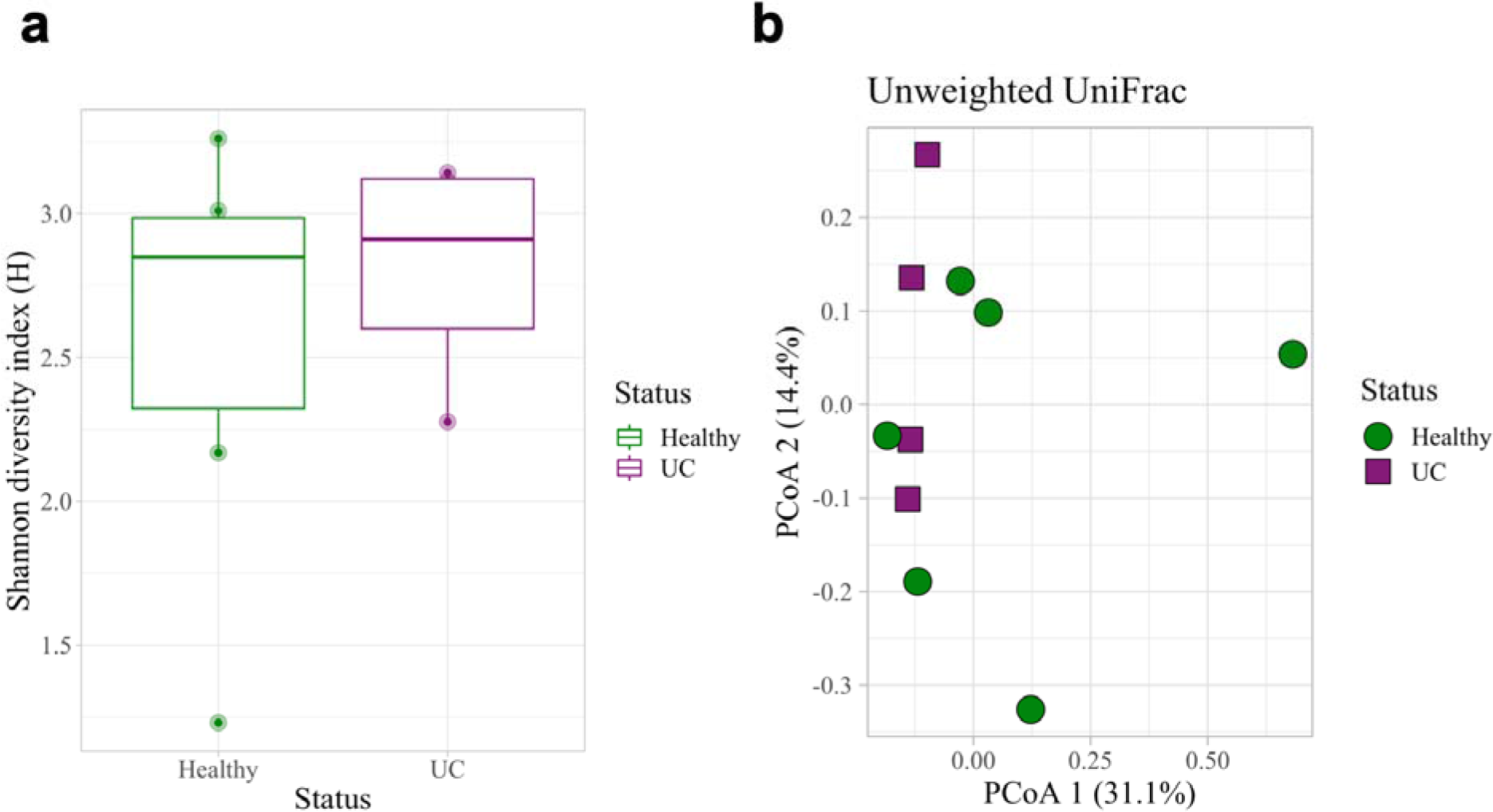
Microbial diversity and composition of fecal samples in dogs with and without UC. (**a**) Fecal microbial diversity did not differ significantly between dogs with and without UC (Kruskal-Wallis, *p* = 0.67). (**b**) Microbial composition also did not differ significantly between healthy dogs and dogs with UC (Unweighted UniFrac, PERMANOVA, *p* = 0.252). Error bars denote standard error.

**Figure 7:**
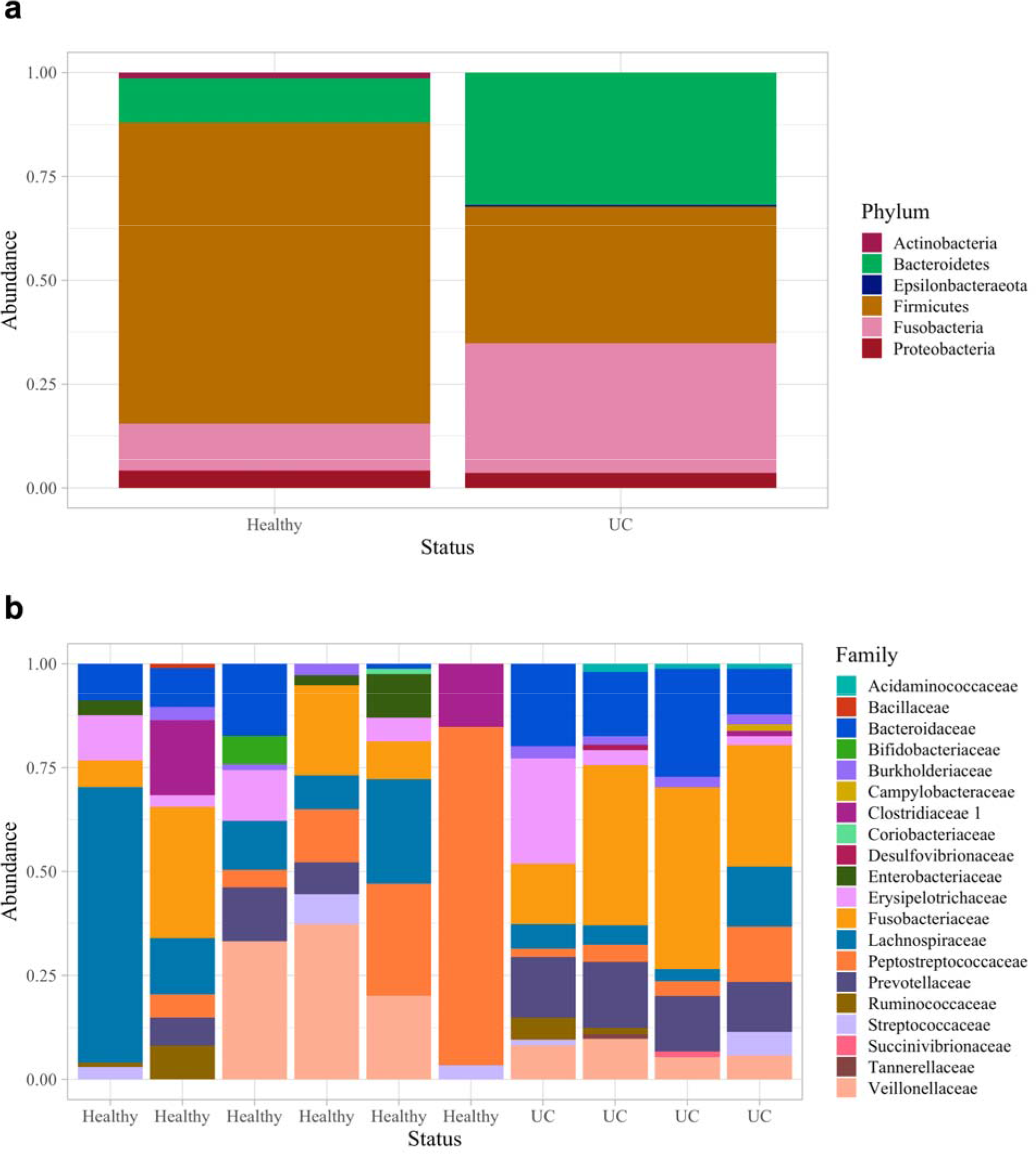
Taxa bar plots of fecal samples in dogs with and without UC. (**a**) Microbial phyla and (**b**) family relative abundances.

To determine how results from this subset of fecal samples compared to a larger sample set, we then analyzed the fecal microbiota of 30 dogs with UC and 30 sex, age, and breed-matched healthy controls (**Supp. Table 4**). Fecal DNA concentrations, 16S reads, and fecal microbial diversity and microbial composition again did not differ significantly between groups (DNA concentration: Wilcoxon Rank Sum test, *p* = 0.515; 16S reads: two-sample t-test, *p* = 0.0697; **Supp. Figure 8; Supp. Table 5**). Firmicutes, Bacteroidetes, and Fusobacteria also remained the most abundant phyla across both groups, and interestingly, Fusobacteriaceae (healthy: 17.4 %; UC: 28 %) and *Fusobacterium* (healthy: 18.5 %; UC: 29.2%) were still the most abundant family and genus in the fecal samples of dogs with UC (**Supp. Figure 9**); although, this difference was still not significant. In fact, no taxa were differentially abundant at the phylum, genus, or ASV levels between groups in the larger sample set (**Supp. Table 5**), suggesting that that *Bacteroides spp.* identified as differentially abundant in the subset was likely an artifact of small sample size.

### 3.3 Microbiota identified in both fecal and urine samples

As the gut can be a source for microbes in the urinary tract [30, 67], we then combined urine and fecal data to determine what ASVs were present in both urine and fecal samples. There were a total of 1,204 ASVs across all urine and fecal samples combined. Sixty-six ASVs were identified in both urine and fecal samples from any dog (**Supp. Table 6**). The most common taxa found in both urine and fecal samples included taxa in the genera *Streptococcus* and *Blautia*. Notably, *Fusobacterium spp., Porphyromonas spp., Campylobacter spp., Helicobacter spp.,* and *Clostridiodes difficile* were also found in both urine and fecal samples. Further, nine ASVs were identified in urine and fecal samples from the same dogs (**Supp. Table 7**). These ASVs included two *Escherichia* or *Shigella spp*., two *Streptococcus spp*., a *Clostridium sensu stricto 1 spp*., *Actinomyces coleocanis*, *Streptococcus minor*, an *Enterococcus spp*., and an uncultured *Peptoclostridium spp*.

## 4. Discussion

The purpose of our study was to characterize the urine and fecal microbiota in a naturally-occurring canine model of UC. We report a decreased urine microbial diversity and altered urine microbial composition in dogs with UC compared to healthy controls. We did not detect significant differences in fecal microbiota between dogs with and without UC; although, *Fusobacterium* was increased in dogs with UC. These results provide a foundation for further exploring the role of microbes in UC in a highly relevant animal model.

### Urine and fecal microbiota associated with UC

The higher concentrations of DNA found in urine from dogs with UC is likely host DNA from epithelial or tumor cells being sloughed into the urine. Notably, urine microbial read numbers did not differ significantly between dogs with and without UC indicating similar amplicon sequencing depths despite differences in DNA concentrations. (Notably, efforts to remove host DNA from UC urine samples prior to sequencing may be beneficial in future microbiome studies employing shotgun metagenomics to ensure that the run is not overwhelmed with host sequences.)

Besides DNA concentrations, we also observed significant differences in urine microbial diversity (Shannon) and composition (Unweighted UniFrac) between dogs with and without UC. In this study, urine microbial diversity was greater in healthy dogs as compared to dogs with UC, a finding that aligns with several studies on urine microbiota in humans with UC [37, 39].

However, there are also studies in humans that report no differences in microbial diversity or decreased diversity in urine from healthy individuals as compared to those with UC [17,35,36,38,42,44,79]. Differences in microbial composition (Unweighted UniFrac) have also been reported in previous human studies on UC [36,38,43,44]. In this study, the four most abundant phyla in urine were Firmicutes, Actinobacteria, Bacteroides, and Proteobacteria. These phyla also dominate the urine microbiota in humans [17,36,38,40,44,45] and have been reported in previous studies on healthy dog urine [80, 81]. In humans, taxa associated with UC vary widely across studies, but *Acinetobacter* and *Actinomyces* have been found at increased abundances in patients with UC across at least three studies [35,42,44]. In this study, we did not see *Acinetobacter* or *Actinomyces spp.* increased in relation to UC, which may be due to small sample sizes and reduced power to detect differentially abundant taxa, or differences between human and canine urine microbiota, or lack of a true link between these taxa and UC.

In relation to fecal microbiota, we did not observe any significant differences in dogs with and without UC. However, intriguingly, *Fusobacterium* was increased in relative abundance (although not significantly) in urine and fecal samples of dogs with UC. One previous study on bladder cancer also reported increased *Fusobacterium* in the urine of individuals (human) with UC [38]. Importantly, taxa in the phyla Fusobacteria are considered normal inhabitants of the canine gastrointestinal tract [82]; although, they are more typically associated with disease in humans. Studies in colorectal cancer have demonstrated direct links between Fusobacteria (*Fusobacterium nucleatum)*and carcinogenesis. Specifically, *Fusobacterium nucleatum*Fap2 protein can bind to host factor Gal-GalNAc which is overexpressed on tumor cells [83] - thereby localizing to tumors where Fap2 can impair host anti-tumor immunity [83]. *Fusobacterium nucleatum* can also induce the host Wnt / beta-catenin pathway resulting in upregulated host cellular proliferation [84]. Future studies are needed to elucidate the potential role of *Fusobacterium* in bladder cancer.

### Microbiota present in both urine and fecal samples

Communication and migration of microbes between the gut and bladder can increase a host’s risk of UTIs and bacteriuria [30]. Microbes may migrate and ascend into the urogenital tract externally from the rectum / anus, or internally via the blood stream [85, 86]. In this study, 66 ASVs were shared between urine and fecal samples. Interestingly, ∼ 59 % of those ASVs (39 / 66) are likely spore-formers (Bacilli, Clostridia, Negativicutes) suggesting that spore formation may more readily enable exchange of microbes between body niches [87, 88]. Among the microbes (ASVs) found in both urine and fecal samples, there were multiple potentially pathogenic taxa: *Campylobacter spp., Helicobacter canis, Clostridiodes difficile, Clostridium baratii, Escherichia* / *Shigella spp., and Enterococcus spp*. There were also a few taxa that have been associated with tumors or directly linked with tumor development or progression in gastrointestinal, oral, and genital cancers: *Fusobacterium spp.* and *Porphyromonas spp.* [89–94]. The shared presence of two *Fusobacterium* ASVs between urine and fecal samples is particularly of interest given the role of *Fusobacterium* in colorectal cancer.

This pilot study is a novel investigation of urine and fecal microbiota in a canine model of UC. The dominant microbial taxa identified in canine urine and fecal samples were similar to those reported in humans. Also, as in humans, altered microbial diversity and composition were observed in dogs with UC as compared to healthy controls. This supports the idea that the microbiota may play a role in UC development, progression, prognosis, or response to treatment, as has been observed in other cancers. Moreover, *Fusobacterium* was increased – albeit not significantly - in both urine and fecal samples of dogs with UC. *Fusobacterium* ASVs were also shared between urine and fecal samples. Taken together, these results provide support for the use of dogs as a model in UC microbiome studies. Additionally, these findings suggest that future work evaluating the role of *Fusobacterium* in UC, and the gut as a potential source of this *Fusobacterium,* may be warranted.

## Funding and Acknowledgements

We are grateful to all individuals involved in sample collection at Purdue University College of Veterinary Medicine (West Lafayette, IN, USA) and to the dogs and dog owners who participated in this study. We also acknowledge the Ohio Supercomputer Center (Columbus, Ohio, USA, established 1987) for computing resources used in this study.

Funding for this project was provided by the Ohio State University College of Veterinary Medicine Department of Veterinary Preventive Medicine (VLH, RM, CM), the Ohio State University Infectious Diseases Institute (VLH, RM, CM), the Ohio State University College of Veterinary Medicine Canine Funds (VLH, WK, DK), and the Ohio State University College of Public Health Collaborative Postdoctoral Research Program Award (MVE).

## Author Contributions

**Conceptualization:** Vanessa L. Hale, Deborah W. Knapp, and William C. Kisseberth **Clinical sample collection, clinical care / monitoring, clinical data extraction:** Deborah W. Knapp, Deepika Dhawan, and William C. Kisseberth

**DNA extraction:** Chris Madden, Ryan Mrofchak, and Morgan V. Evans

**Data processing, analysis:** Ryan Mrofchak, Morgan, V. Evans, Chris Madden, and Deborah W. Knapp

**Data interpretation and conclusions:** Ryan Mrofchak, Vanessa L. Hale, Chris Madden, and Deborah W. Knapp

**Manuscript writing:** Ryan Mrofchak, Vanessa L. Hale, and Chris Madden

**Manuscript editing:** Chris Madden, Deepika Dhawan, Deborah W. Knapp, and William C. Kisseberth

## Supplemental Material

**Supplemental Table 1:**
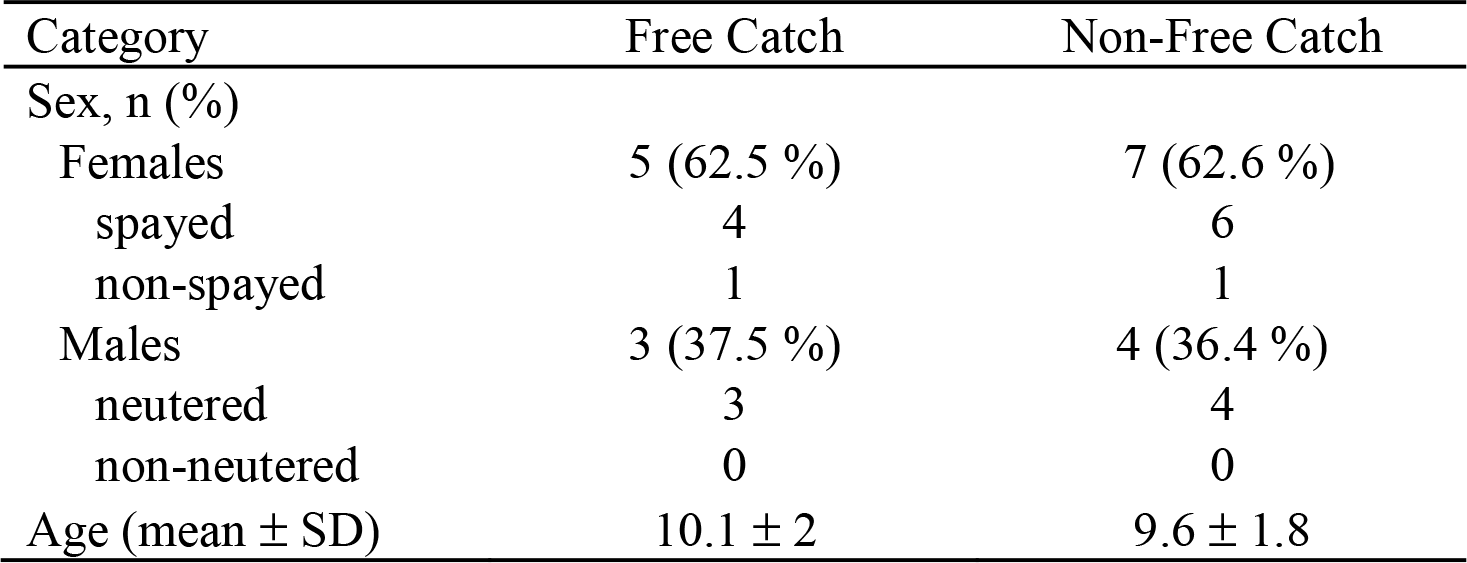
Demographics of dogs with urine samples collected via free catch and non-free catch methods. All dogs had urothelial carcinoma. Eight dogs had urine collected via mid-stream free catch while eleven dogs were sampled via non-free catch methods including cystoscopy or catheterization.

**Supplemental Table 2:** Dominant taxa in urine from dogs with UC by collection method. Relative abundance of the top three taxa in free catch and non-free catch urine at the phylum and genera levels. All urine was collected from dogs with UC.

| Free Catch Urine |  | Non-free Catch Urine |  |
| --- | --- | --- | --- |
| Phylum |  |  |  |
| Firmicutes | 70.3 % | Firmicutes | 33 % |
| Proteobacteria | 20.1 % | Tenericutes | 26.7 % |
| Bacteroidetes | 5.98 % | Proteobacteria | 26.7 % |
| Genera |  |  |  |
| *Staphylococcus* | 43.2 % | *Mycoplasma* | 18.3 % |
| *Streptococcus* | 12.6 % | *Escherichia-Shigella* | 18.1 % |
| *Pantoea* | 11.4 % | *Enterococcus* | 9.73 % |

**Supplemental Table 3:** Contaminant ASVs. Using the frequency and prevalence methods (threshold value of 0.5) in the R package decontam v.1.10.0, putative contaminant ASVs were identified and bioinformatically removed prior to further analyses.

| Putative urine contaminants (ASVs) |
| --- |
| D_1__Tenericutes;D_2__Mollicutes RF39;D_4__uncultured prokaryote;D_5__uncultured prokaryote;D_6__uncultured prokaryote |
| D_1__Deinococcus-Thermus;D_2__Deinococci;D_3__Thermales;D_4__Thermaceae;D_5__Thermus |
| D_1__Actinobacteria;D_2__Actinobacteria;D_3__Micrococcales;D_4__Micrococcaceae;D_5__Micrococcus |
| D_1__Proteobacteria;D_2__Gammaproteobacteria;D_3__Betaproteobacteriales;D_4__Burkholderiaceae;D_5__Cupriavidus |
| D_1__Proteobacteria;D_2__Gammaproteobacteria;D_3__Betaproteobacteriales;D_4__Burkholderiaceae |
| D_1__Bacteroidetes;D_2__Bacteroidia;D_3__Bacteroidales;D_4__Prevotellaceae;D_5__Prevotella9;D_6__uncultured bacterium |
| D_1__Kiritimatiellaeota;D_2__Kiritimatiellae;D_3__WCHB1-41;D_4__uncultured rumen bacterium;D_5__uncultured rumen bacterium;D_6__uncultured rumen bacterium |
| D_1__Bacteroidetes;D_2__Bacteroidia;D_3__Bacteroidales;D_4__Prevotellaceae |
| D_1__Firmicutes;D_2__Bacilli;D_3__Lactobacillales;D_4__Lactobacillaceae;D_5__Lactobacillus;D_6__Lactobacillus iners AB-1 |
| D_1__Firmicutes;D_2__Bacilli;D_3__Lactobacillales;D_4__Lactobacillaceae;D_5__Cytophaga |
| D_1__Verrucomicrobia;D_2__Verrucomicrobiae;D_3__Opitutaceae;D_4__Opitutaceae;D_5__Lacunisphaera;D_6__Opitutus sp. WS3(2011) |
| D_1__Bacteroidetes;D_2__Bacteroidia;D_3__Bacteroidales;D_4__Prevotellaceae;D_5__Prevotella 9 |
| D_1__Proteobacteria;D_2__Alphaproteobacteria;D_3__Rhizobiales;D_4__Xanthobacteraceae;D_5__Bradyrhizobium |
| Putative fecal contaminants (ASVs) |
| D_0__Bacteria |
| D_1__Firmicutes;D_2__Negativicutes;D_3__Selenomonadales;D_4__Veillonellaceae;D_5__Veillonella |
| D_1__Firmicutes;D_2__Bacteroidia;D_3__Bacteroidales;D_4__Prevotellaceae;D_5__Prevotella 9 |
| D_1__Firmicutes;D_2__Bacilli;D_3__Bacillales;D_4__Staphylococcaceae;D_5__Staphylococcus |
| D_1__Actinobacteria;D_2__Coriobacteriia;D_3__Coriobacteriales;D_4__Atopobiaceae;D_5__Coriobacteriaceae UCG-002 |
| D_1__Proteobacteria;D_2__Gammaproteobacteria;D_3__Enterobacteriales;D_4__Enterobacteriaceae |
| D_1__Actinobacteria;D_2__Coriobacteriia;D_3__Coriobacteriales;D_4__Atopobiaceae;D_5__Coriobacteriaceae UCG-002 |

**Supplemental Table 4:** Demographics of larger canine cohort from which fecal samples were collected. Fecal samples were collected from dogs with UC (n = 30) and age-, sex-, breed-matched healthy controls (n = 30).

| Category | Healthy | UC |
| --- | --- | --- |
| Sex, n (%) |  |  |
| Females | 16 (53.3 %) | 16 (53.3 %) |
| spayed | 15 | 15 |
| non-spayed | 1 | 1 |
| Males | 14 (46.7 %) | 14 (46.7 %) |
| neutered | 11 | 11 |
| non-neutered | 3 | 3 |
| Age (mean ± SD) | 10 ± 1.76 | 10.4 ± 1.97 |

**Supplemental Table 5.** Microbial diversity and composition of fecal samples from healthy dogs and dogs with UC. There were no significance differences in microbial diversity or composition between dogs with UC (n = 30) and sex-, age-, and breed-matched healthy controls (n = 30). ANCOM – Analysis of Composition of Microbiome.

|  | **Metric** | **Fecal samples from healthy dogs vs. dogs with UC** |
| --- | --- | --- |
| **Alpha Diversity** | Shannon Diversity Index<br>Kruskal-Wallis | $p = 0.214$ |
| | Simpson Diversity Index<br>Kruskal-Wallis | $p = 0.506$ |
| | Observed Features<br>Kruskal-Wallis | $p = 0.336$ |
| **Beta Diversity** | Bray Curtis<br>PERMANOVA | $p = 0.468$ |
| | UnWeighted UniFrac<br>PERMANOVA | $p = 0.134$ |
| | Weighted UniFrac<br>PERMANOVA | $p = 0.0819$ |
| **Differentially Abundant Taxa** | Phylum<br>ANCOM | No differentially abundant taxa |
|  | Genus<br>ANCOM | No differentially abundant taxa |
|  | ASV<br>ANCOM | No differentially abundant taxa |

**Supplemental Table 6:**
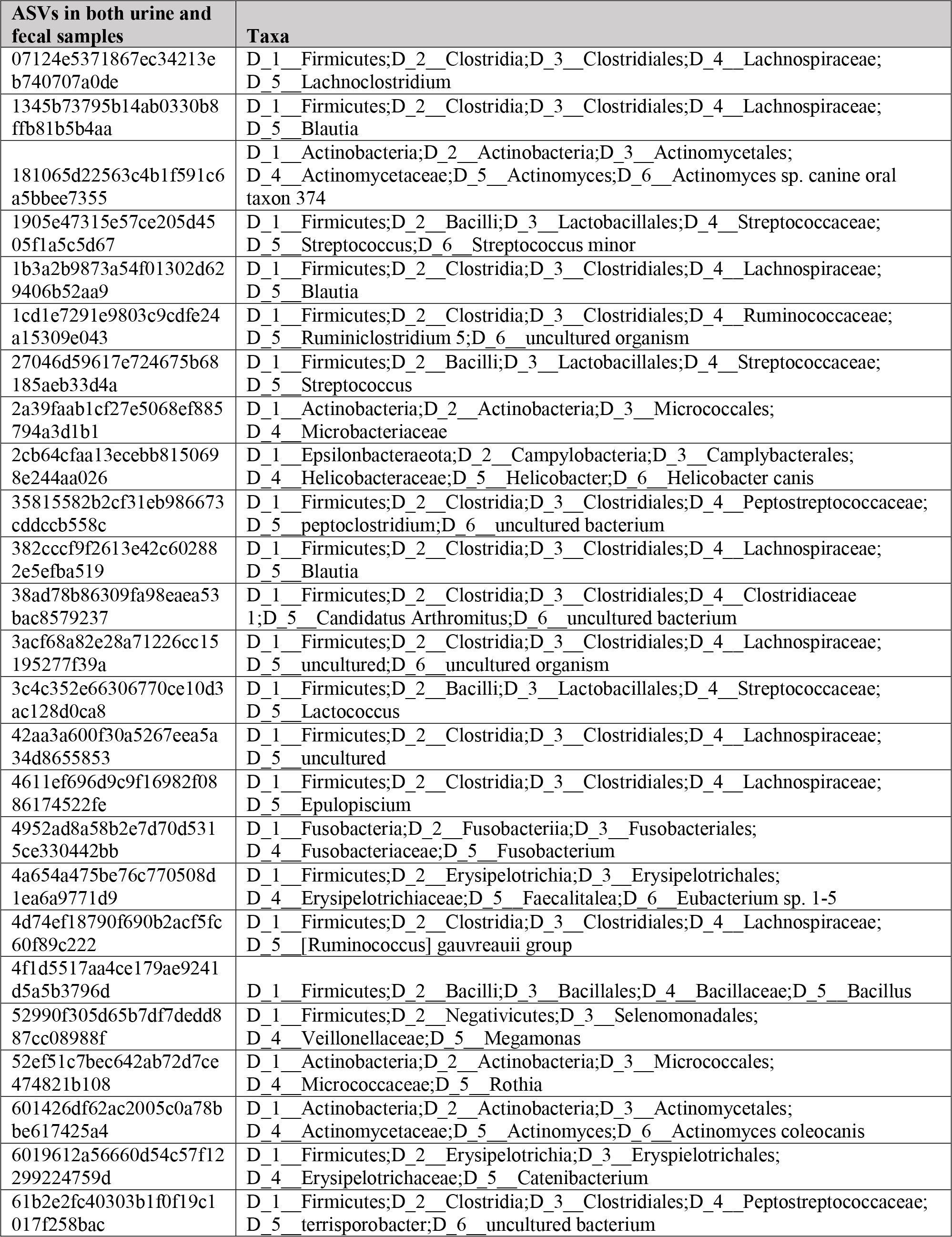
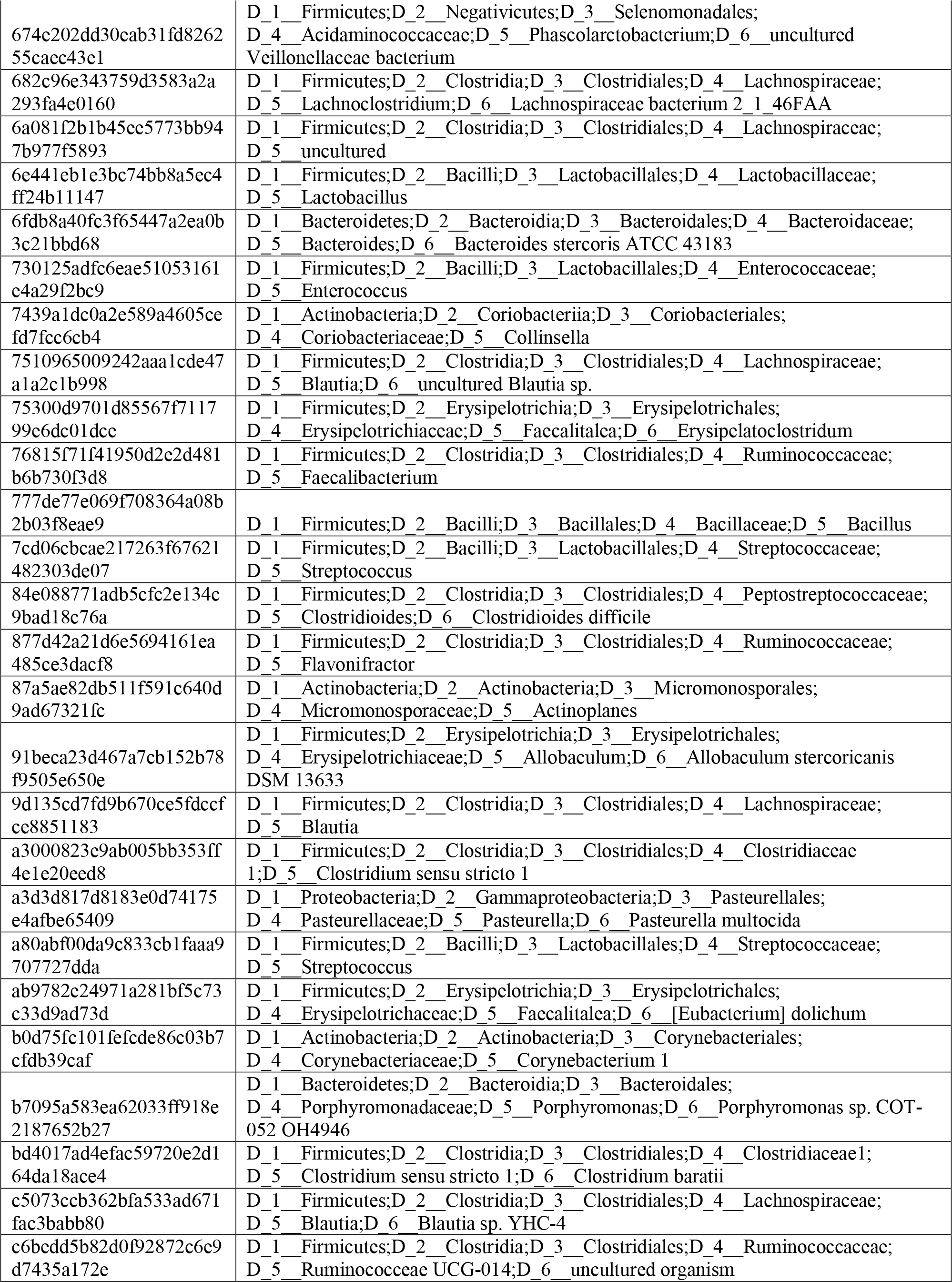
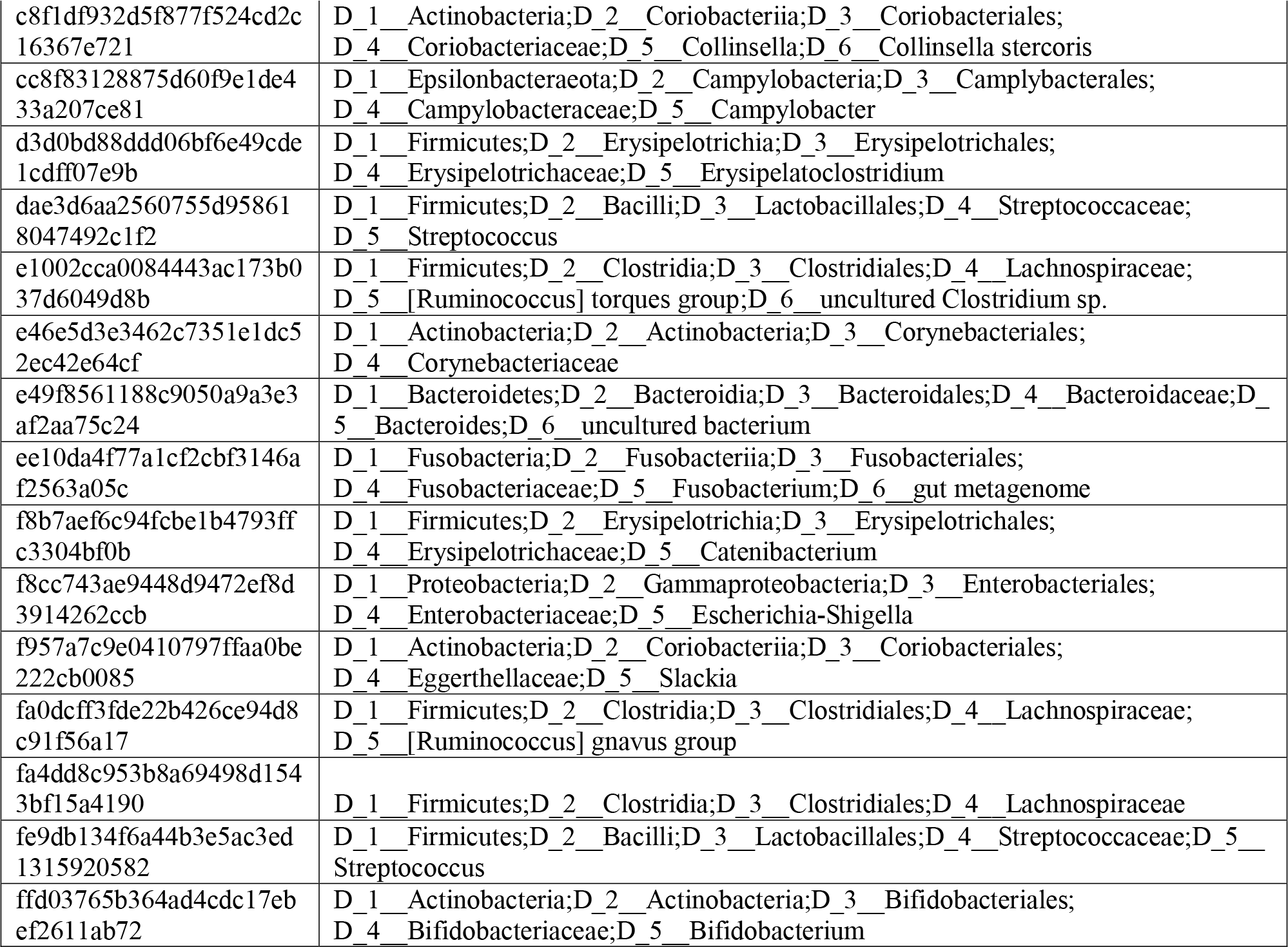
ASVs identified in both urine and fecal samples. There were 66 ASVs found in both urine and fecal samples of any dog.

**Supplemental Table 7:** ASVs in urine and fecal samples from the same dog. Four dogs contained ASVs that were found in both their urine and fecal samples.

| ASVs in both urine and fecal samples by dog | Taxa |
| --- | --- |
| **Dog 1 - UC** |  |
| f8cc743ae9448d9472ef8d3914262ccb | D_1__Proteobacteria;D_2__Gammaproteobacteria;D_3__Enterobacteriales;D_4__Enterobacteriaceae;D_5__Escherichia-Shigella |
| **Dog 2 - UC** |  |
| 27046d59617e724675b68185aeb33d4a | D_1__Firmicutes;D_2__Bacilli;D_3__Lactobacillales;D_4__Streptococcaceae;D_5__Streptococcus |
| **Dog 3 - Healthy** |  |
| f8cc743ae9448d9472ef8d3914262ccb | D_1__Proteobacteria;D_2__Gammaproteobacteria;D_3__Enterobacteriales;D_4__Enterobacteriaceae;D_5__Escherichia-Shigella |
| 1878459013cf15f2993a81c14978c980 | D_1__Firmicutes;D_2__Bacilli;D_3__Lactobacillales;D_4__Streptococcaceae;D_5__Streptococcus |
| a3000823e9ab005bb353ff4e1e20eed8 | D_1__Firmicutes;D_2__Clostridia;D_3__Clostridiales;D_4__Clostridiales 1;D_5__Clostridium sensu stricto 1 |
| 601426df62ac2005c0a78bbe617425a4 | D_1__Actinobacteria;D_2__Actinobacteria;D_3__Actinomycetales;D_4__Actinomycetaceae;D_5__Actinomyces;D_6__Actinomyces coleocanis |
| 1905e47315e57ce205d4505f1a5c5d67 | D_1__Firmicutes;D_2__Bacilli;D_3__Lactobacillales;D_4__Streptococcaceae;D_5__Streptococcus;D_6__Streptococcus minor |
| **Dog 4 - Healthy** |  |
| 730125adfc6eae51053161e4a29f2bc9 | D_1__Firmicutes;D_2__Bacilli;D_3__Lactobacillales;D_4__Enterococcaceae;D_5__Enterococcus |
| 35815582b2cf31eb986673cddccb558c | D_1__Firmicutes;D_2__Clostridia;D_3__Clostridiales;D_4__Peptostreptococcaceae;D_5__Peptoclostridium;D_6__uncultured bacterium |

**Supplemental Figure 1:**
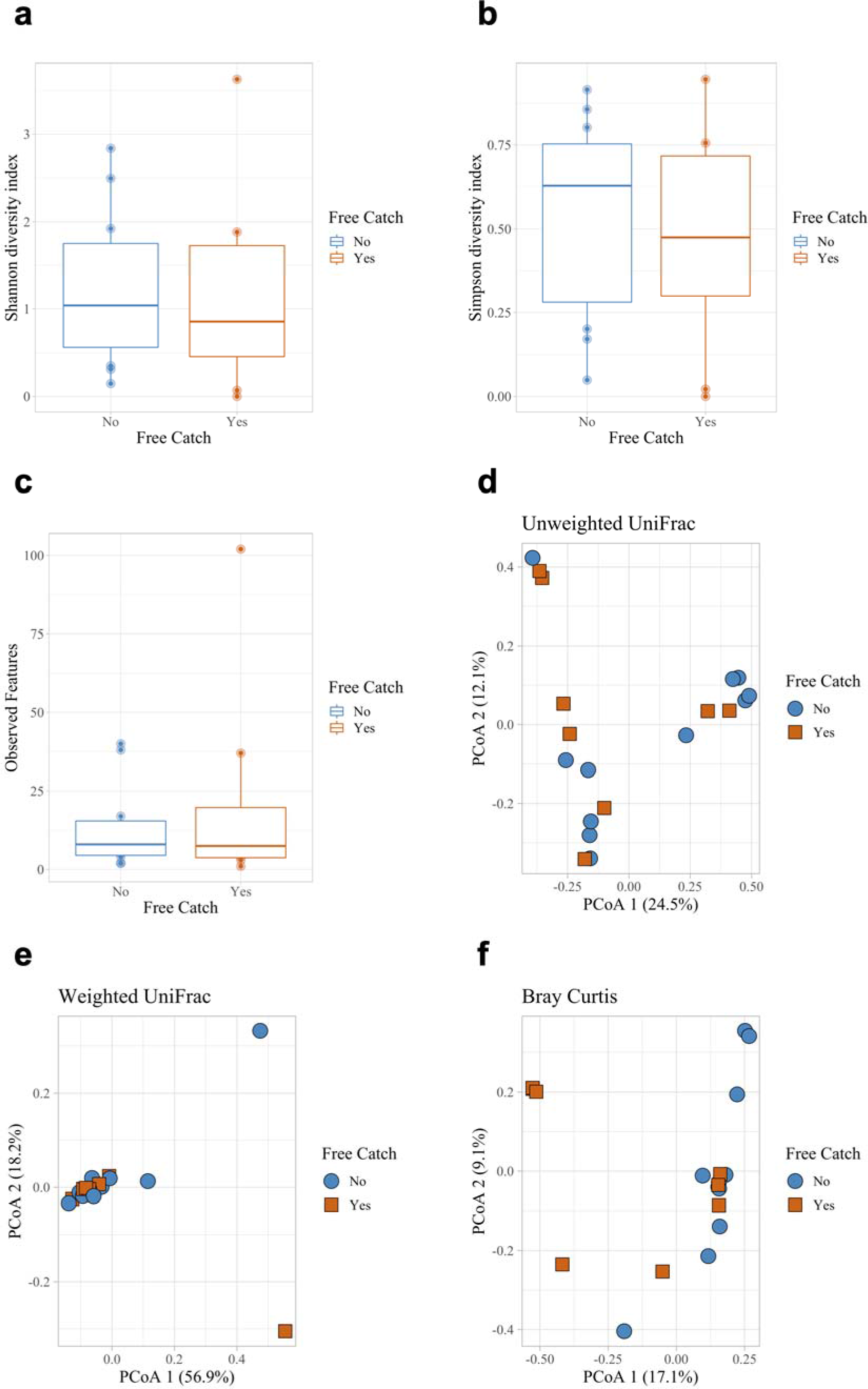
Urine microbial community diversity and composition by collection method in dogs with UC (rarefied data). Dogs with UC were sampled via free catch (n = 8) and non-free catch (n = 11) methods. Samples were rarefied at 1000 reads. There were no significant differences in microbial diversity between collection methods as assessed via (**a**) Shannon (Kruskal-Wallis: *p* = 0.62) or **b**) Simpson diversity indices (*p* = 0.68) or (**c**) Observed Features (richness) (*p* = 0.901). The microbial composition of free-catch urine did not differ significantly from non-free catch urine based on (**d**) Unweighted (PERMANOVA, *p* = 0.328) or (**e**) Weighted UniFrac distance matrices (*p* = 0.485) but did differ significantly based on (**f**) Bray Curtis (*p* = 0.008). Error bars denote standard error.

**Supplemental Figure 2:**
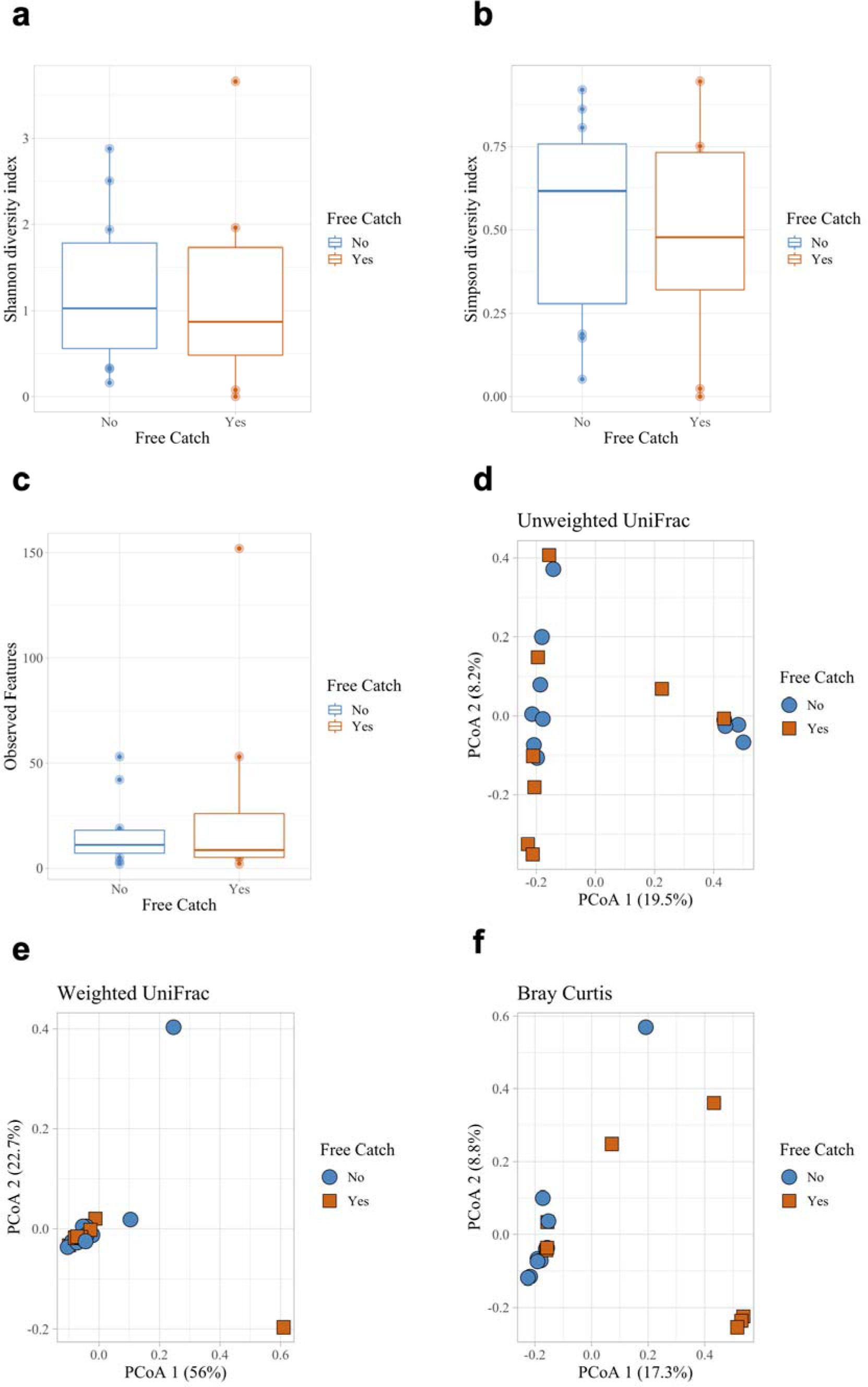
Urine microbial community diversity and composition by collection method in dogs with UC (unrarefied data). Dogs with UC were sampled via free catch (n = 8) and non-free catch (n = 11) methods. Data are non-rarefied. There were no significant differences in alpha diversity between collection methods as assessed using the (**a**) Shannon (Kruskal-Wallis: *p* = 0.68) or **b**) Simpson diversity indices (*p* = 0.68) or (**c**) Observed Features (richness) (*p* = 0.901). The microbial composition of free-catch urine did not differ significantly from non-free catch urine based on (**d**) Unweighted (PERMANOVA, *p* = 0.342) or (**e**) Weighted UniFrac distance matrices (*p* = 0.54) but did differ significantly based on (**f**) Bray Curtis (*p* = 0.005). Error bars denote standard error.

**Supplemental Figure 3:**
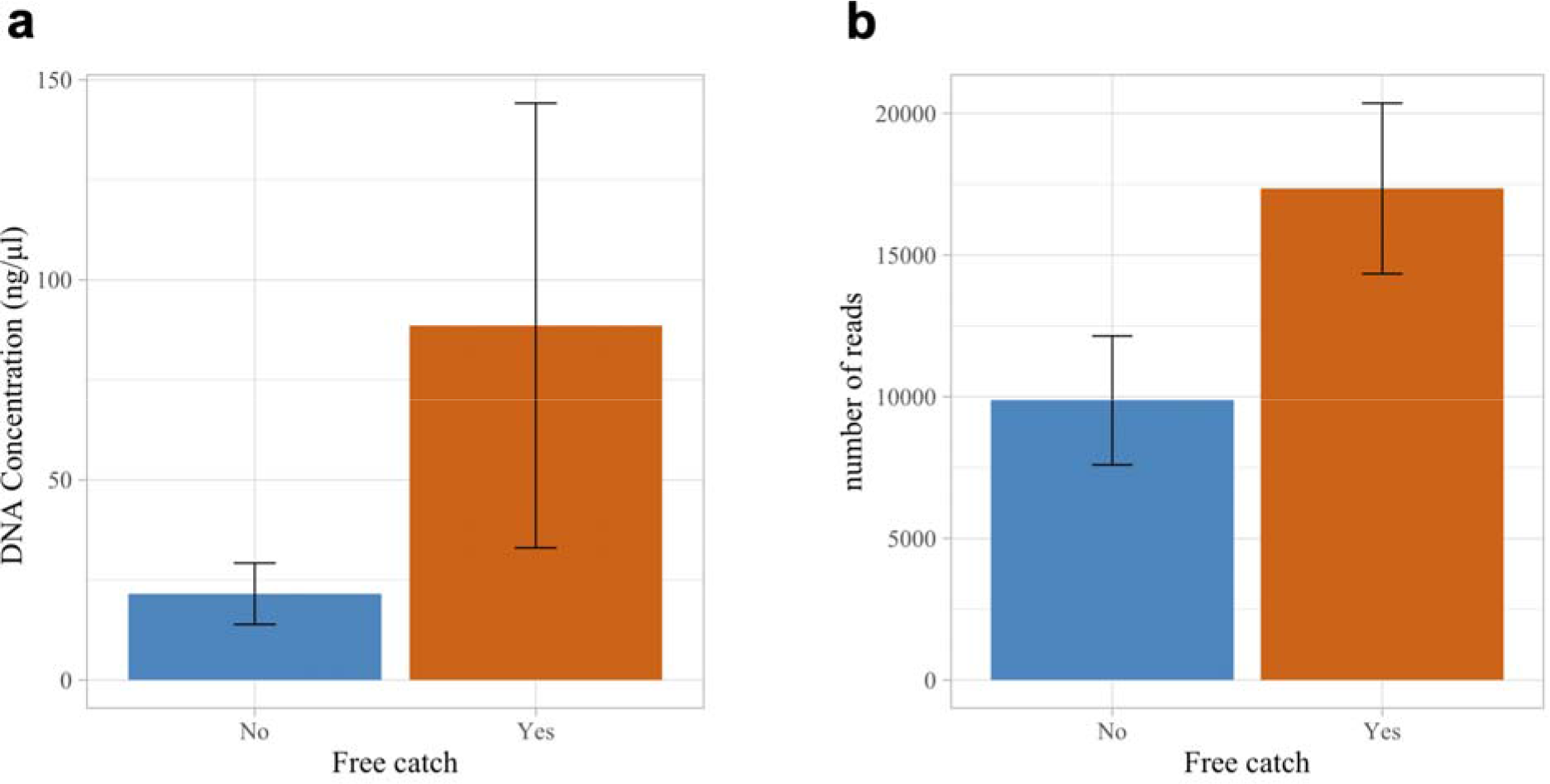
DNA Concentrations and 16S reads by urine collection method. (**a**) Urine DNA concentrations and (**b**) 16S reads in dogs with UC sampled via free catch or non-free catch methods (cystoscopy, catheterization). DNA concentrations and 16S reads were greater, although not significantly, in mid-stream free catch urine samples (DNA concentration: Wilcoxon Test, *p* = 0.778; 16S reads: two-sample t-test, *p* = 0.067). Error bars denote standard error.

**Supplemental Figure 4:**
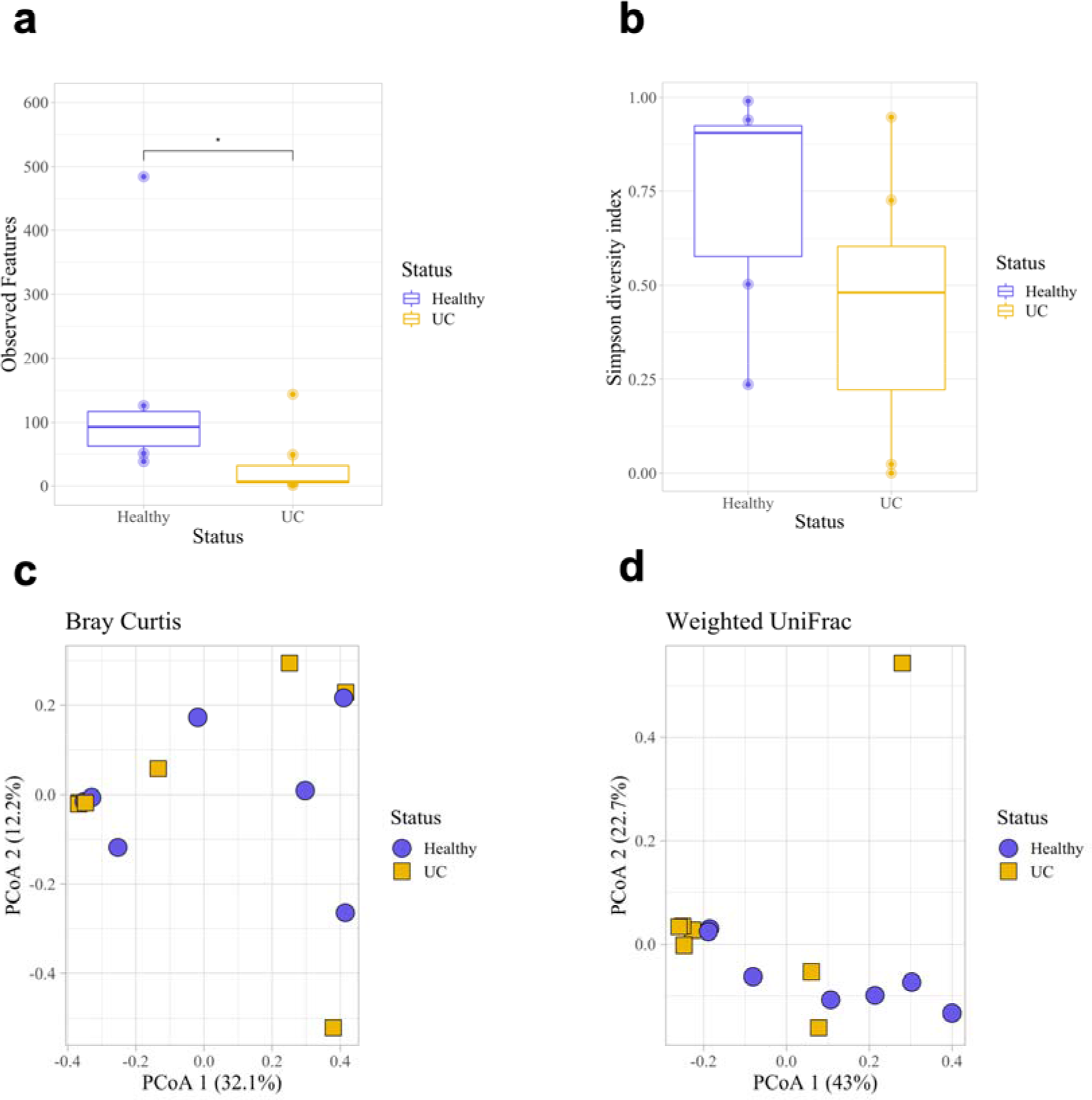
Urine microbial diversity and composition in dogs with and without UC. Dogs with UC had lower microbial diversity compared to healthy dogs based on (**a**) Observed Features (richness) and the (**b**) Simpson diversity index; however, only Observed Features was statistically significant (Kruskal-Wallis: Observed Features, *p* = 0.025; Simpson, *p* = 0.133). Microbial composition did not differ significantly based on (**c**) Bray Curtis or (**d**) Weighted UniFrac distance matrices (PERMANOVA: Bray Curtis, *p* = 0.888; Weighted UniFrac, *p* = 0.168). Error bars denote standard error. Statistical significance is represented by stars: * < 0.05, ** < 0.001, *** < 0.0001

**Supplemental Figure 5:**
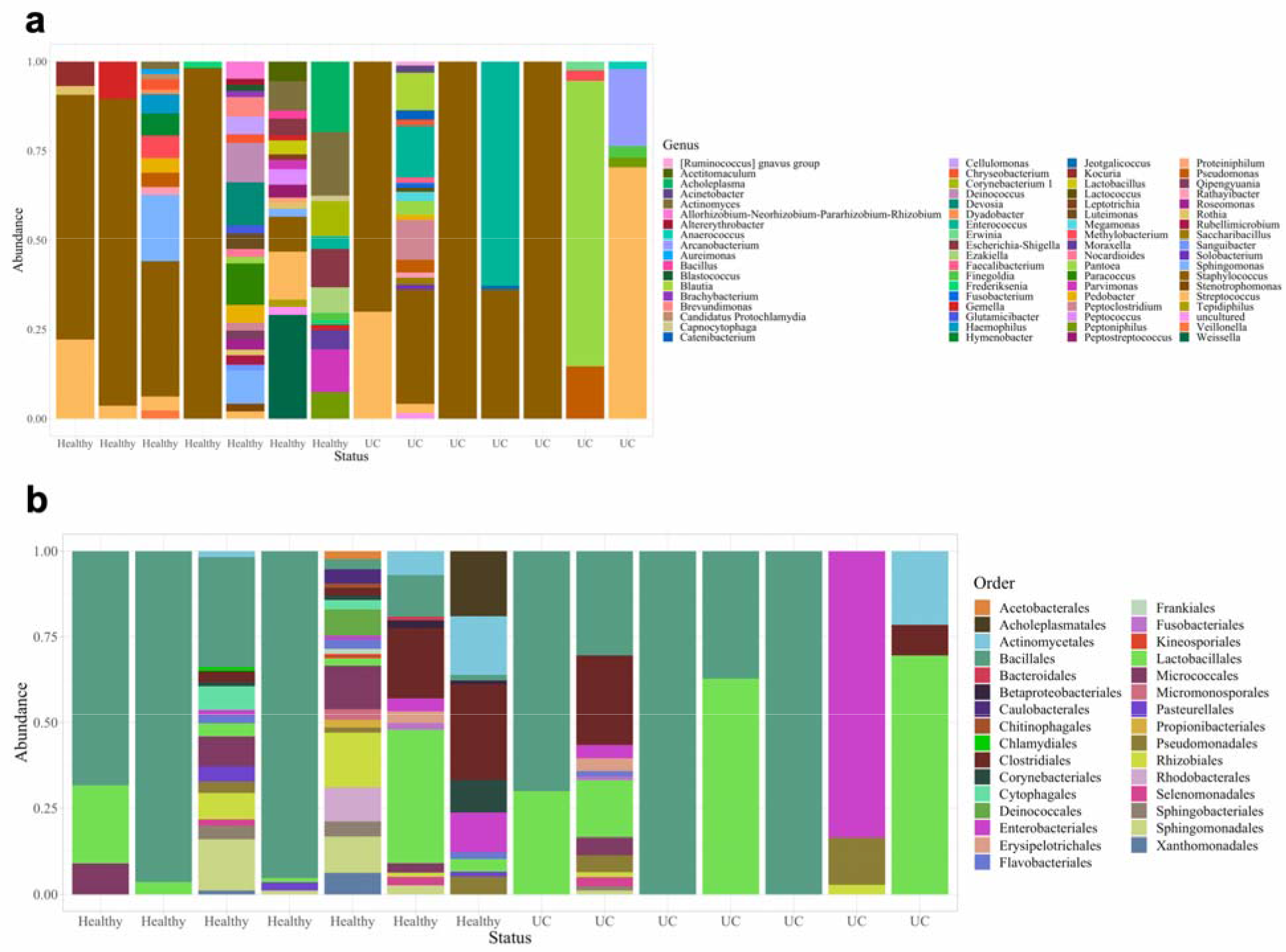
Taxa bar plots of urine samples in dogs with and without UC. (**a**) Microbial genera and (**b**) order relative abundances.

**Supplemental Figure 6:**
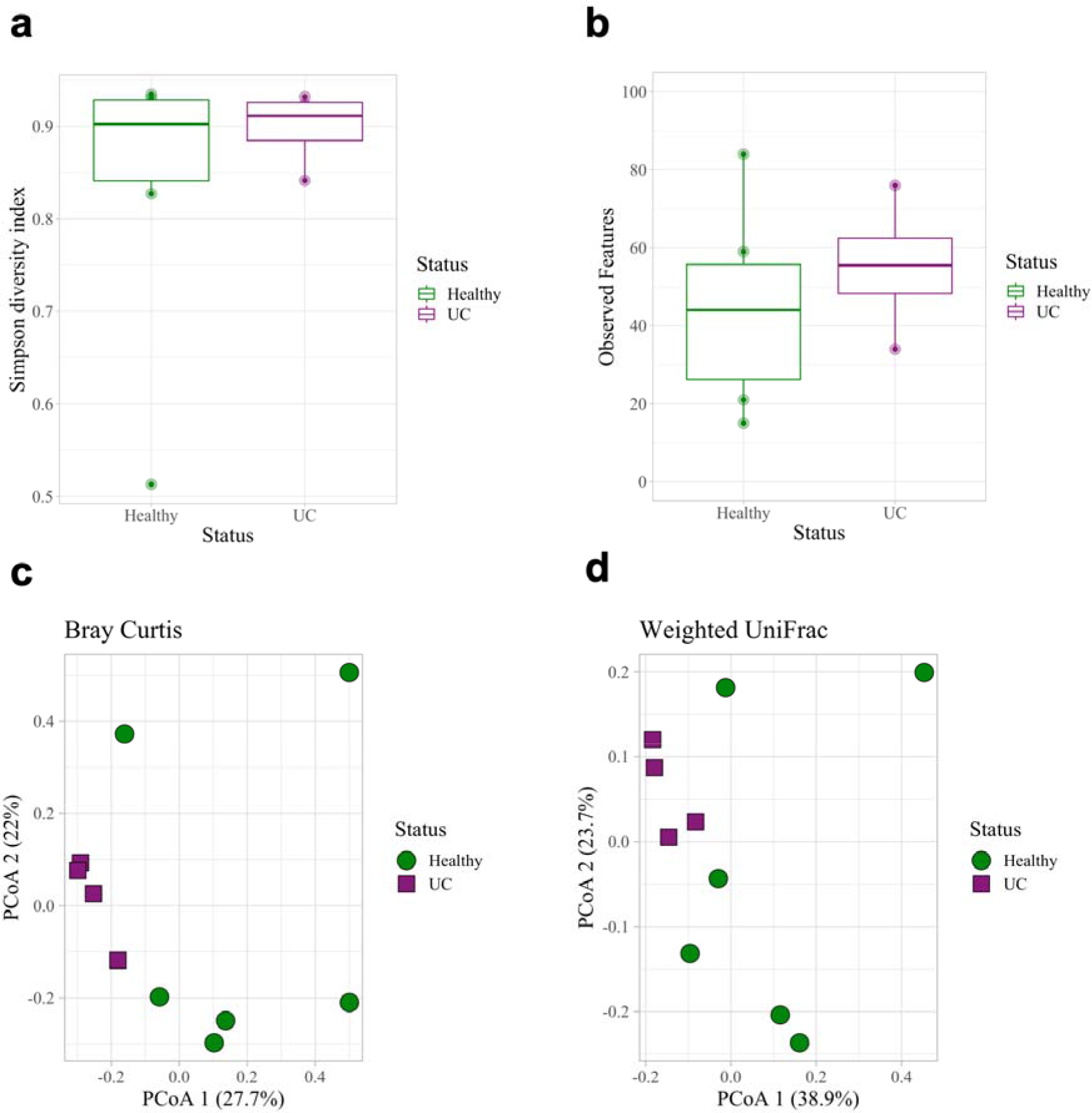
Fecal microbial diversity and composition in dogs with and without UC. Fecal microbial diversity did not differ significantly in dogs with (n=4) or without (n=6) UC based on (**a**) Observed Features (richness) and the (**b**) Simpson diversity index (Kruskal-Wallis: Observed Features, *p* = 0.67; Simpson, *p* = 0.522). Microbial composition also did not differ significantly based on (**c**) Bray Curtis or (**d**) Weighted UniFrac distance matrices (PERMANOVA: Bray Curtis, *p* = 0.06; Weighted UniFrac, *p* = 0.06). Error bars denote standard error.

**Supplemental Figure 7:**
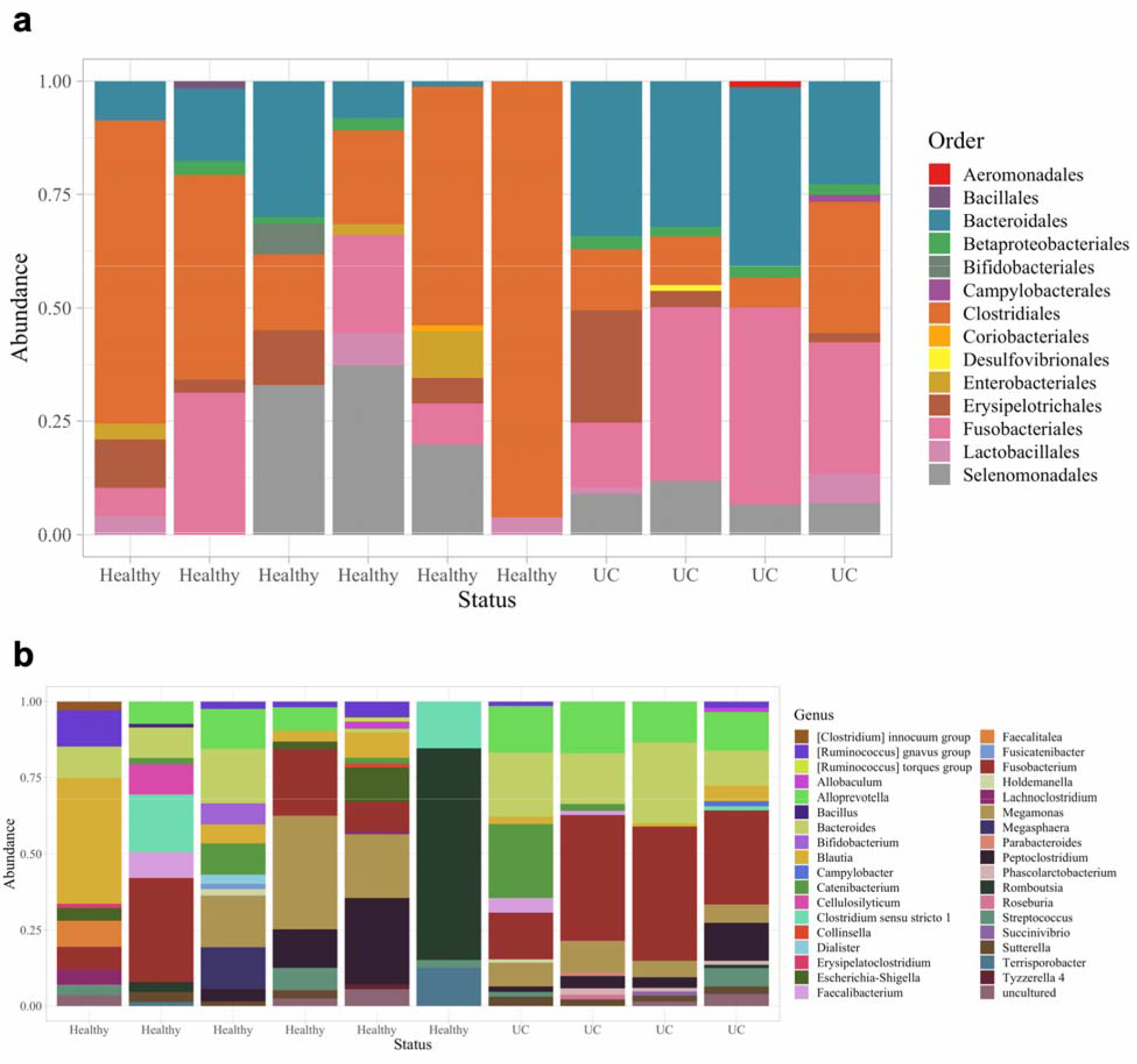
Taxa bar plots of fecal samples. (**a**) Microbial order and (**b**) genera relative abundances in dogs with (n=4) and without UC (n=6).

**Supplemental Figure 8:**
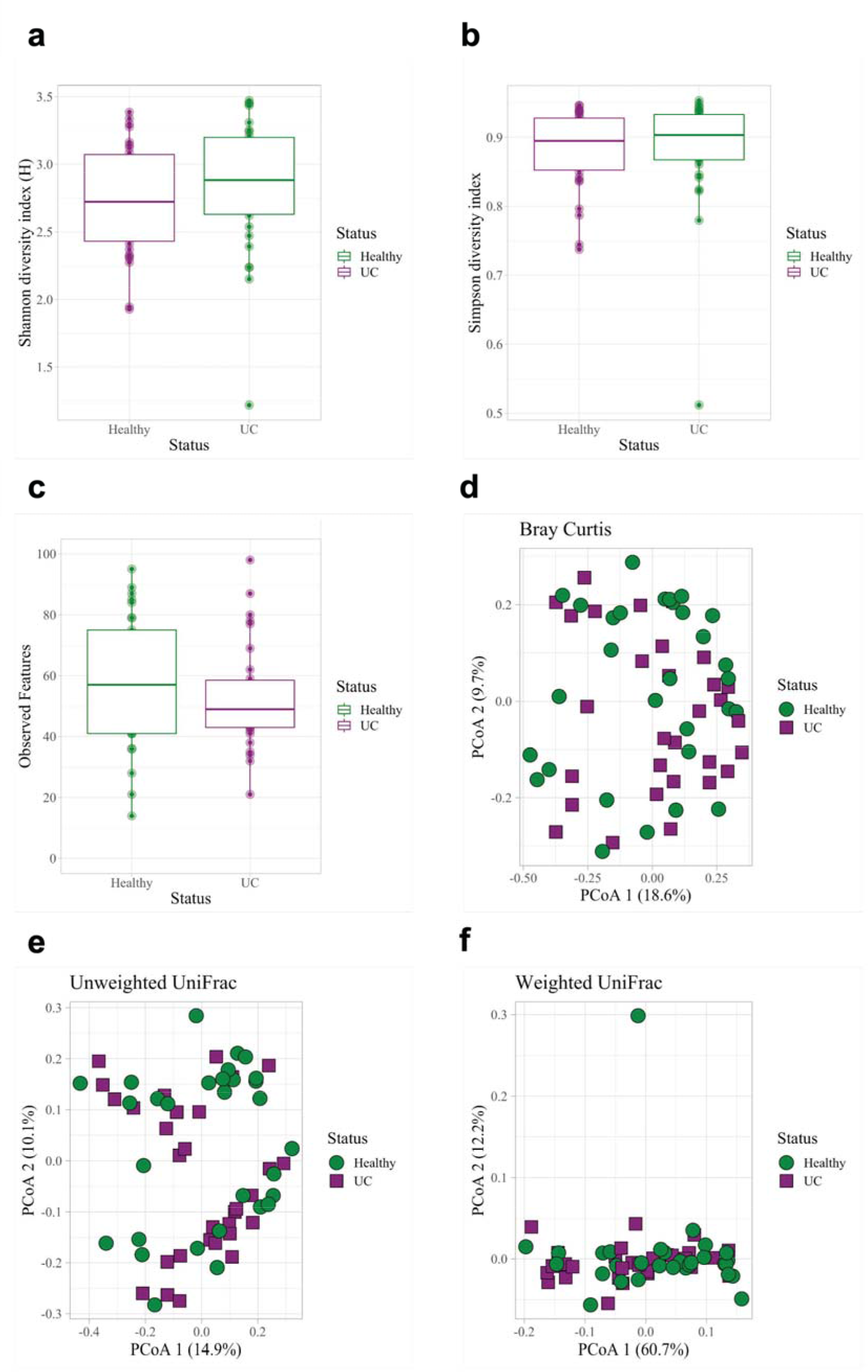
Fecal microbial diversity and composition. We compared fecal microbiota in dogs with UC (n = 30) and sex-, age-, and breed-matched healthy controls (n = 30). There were no significant differences in microbial diversity by (**a**) Shannon (Kruskal-Wallis, *p* = 0.214), (**b**) Simpson (Kruskal-Wallis, *p* = 0.506), or (**c**) Observed Features (Kruskal-Wallis, *p* = 0.336). There were also no significant differences in microbial composition by (**d**) Bray Curtis (PERMANOVA, *p* = 0.468), (**e**) Unweighted UniFrac (PERMANOVA, *p* = 0.134), or (**f**) Weighted UniFrac distance matrices (PERMANOVA, *p* = 0.0819).

**Supplemental Figure 9:**
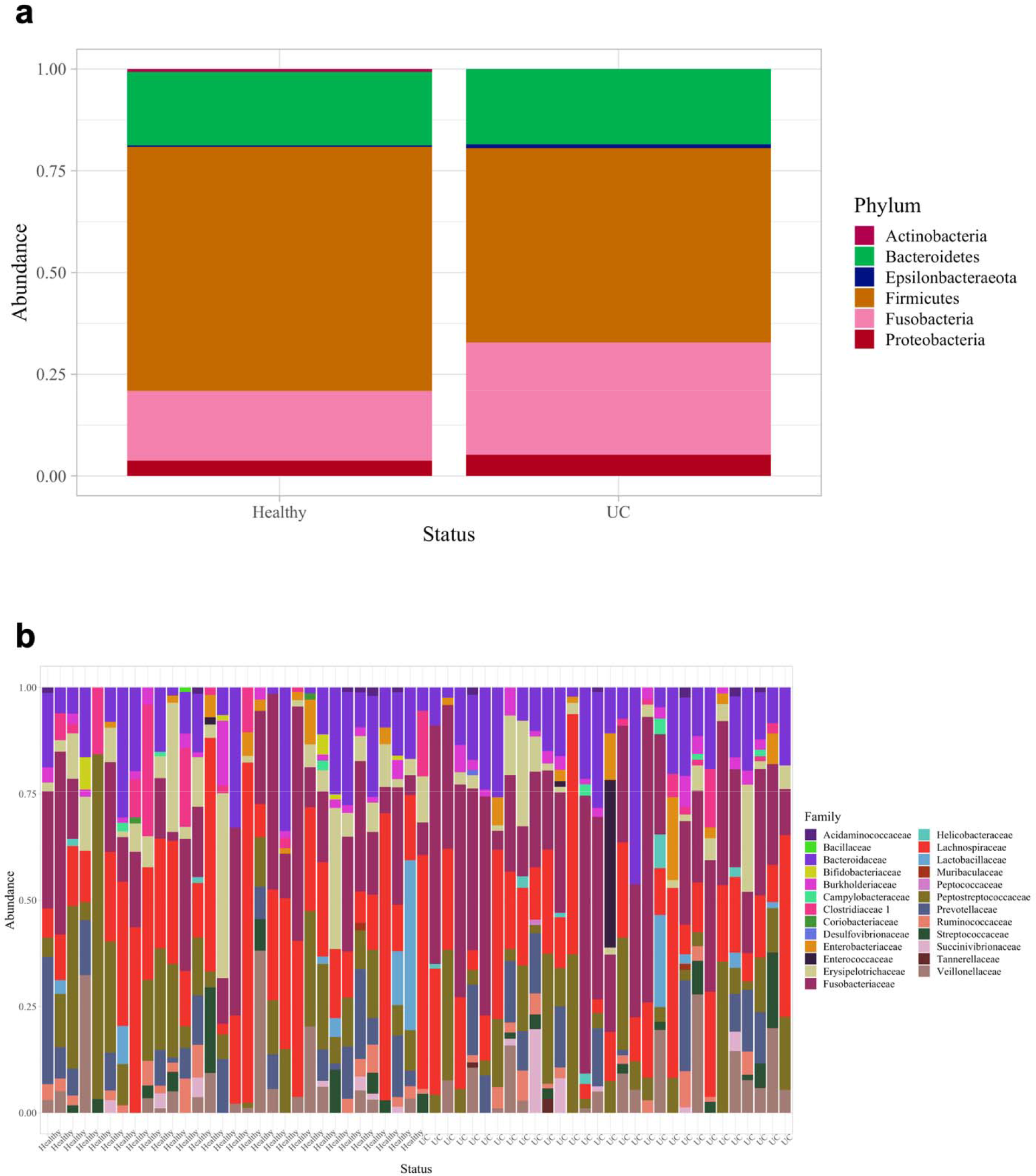
Fecal microbial taxa bar plots. Relative abundances of fecal microbiota at the (**a**) phyla and (**b**) family levels from dogs with UC (n = 30) and age-, sex-, and breed-matched healthy controls (n = 30).

